# Transcriptomic Analysis of the Effect of Remote Ischaemic Conditioning in an Animal Model of Necrotising Enterocolitis

**DOI:** 10.1101/2023.10.24.563747

**Authors:** Ian Jones, Jane Collins, Nigel Hall, Ashley Heinson

**Author notes:** Data Availability The datasets generated and/or analysed during the current study are available in the University of Southampton repository, https://eprints.soton.ac.uk/475296/.

## Abstract

**Background and Aims:** Previously, we reported that remote ischaemic conditioning (RIC) reduces bowel injury in an animal model of Necrotising enterocolitis (NEC). We investigated the mechanisms by which RIC confers this protective effect using RNA-Seq.

**Methods:** Related rat-pups were randomly assigned to four groups: SHAM, intestinal ischaemia-reperfusion injury (IRI), RIC and RIC+IRI. Anaeasthetised IRI animals underwent 40 minutes of intestinal ischaemia, followed by 90 minutes of reperfusion. Animals that underwent RIC had three 5 minute cycles of alternating ischaemia/reperfusion by ligature application to the hind limb.

Illumina NextSeq 550 High Throughput NG Sequencing and genome alignment was performed with Qiagen’s CLC read mapper to produce raw gene counts. Transcriptome analysis was done using *R* v 3.6.1.

**Results:** Differential expression testing showed 868 differentially expressed genes, in animals exposed to RIC alone compared to SHAM, 135 differentially expressed with IRI/RIC compared to IRI alone. Comparison between these two sets showed 25 genes were differentially expressed in both groups. Of these, several genes involved in pro-inflammatory pathways, including NF-ĸβ2, Cxcl1, SOD2 and Map3k8, all showed reduced expression in response to RIC. Targeted analysis revealed increased expression in PI3K which is part of the RISK-pathway identified as a response to RIC in cardiac tissue.

**Conclusions:** Expression patterns suggest that within the intestine, RIC suppresses pro-inflammatory pathways and that an equivalent of the RISK-pathway may be present in the intestine. The cross-over between the pro-inflammatory pathways suppressed here and those that are involved in several stages of the pathogenesis of NEC, further support the potential for RIC as a treatment for NEC.

## Introduction

Necrotising enterocolitis (NEC) is a devastating disease with a high mortality and morbidity for the survivors^1^ The pathophysiology of NEC is complex, but ischaemia-reperfusion injury (IRI) is a common end-point of different, overlapping pathophysiological processes.^2^

Remote ischaemic conditioning (RIC) is an endogenous phenomenon whereby the application of short cycles of non-injurious ischaemia and reperfusion to one organ system (i.e. skeletal muscle) provides a systemic protective effect against ischaemia-reperfusion injury.^3^ We have previously reported the protective effect of remote ischaemic pre-conditioning (pre-RIC) in an animal model based on ischaemia-reperfusion injury to the intestine in a rat pup.^4^ In this model, the application of pre-RIC resulted in a dramatic reduction in intestinal injury.

Whole transcriptome analysis of mRNA is a powerful tool for studying complex biochemical pathways. In recent years, a few papers have studied the changes in gene expression triggered by remote ischaemic conditioning (RIC).

Yoon *et al.* (2015)^5^ used microarray analysis to study the gene expression changes in a porcine model of preRIC and renal Ischaemia reperfusion injury (IRI). They found that RIC had effects on the expression of multiple cytokines (including Interleukin-10 (IL-10) and Transforming growth factor beta (TGF-β)), as well as modulating the complement and coagulation cascades. However, there are potentially significant differences in their experimental design. Firstly, in the model of renal IRI, RIC did not alter the primary end-point, which was a surrogate for renal function. Secondly the tissues were harvested two days after the IRI and thus do not capture the immediate changes in gene-expression.

The analysis of gene expression in a porcine model of myocardial infarction with post-ischaemic conditioning was reported by Lukovic *et al.* in 2019.^7^ Unsurprisingly many of the gene expression changes seen in the myocardium are the same in animals exposed to post conditioning as those who were not; both groups undergoing a myocardial injury. However, there were distinct genes whose regulation was different in animals who underwent conditioning. They concluded that ischaemic conditioning downregulates ECM-proteinases, ribosomal subunits and platelet and leukocyte adhesion molecules. Additionally, post-conditioning inhibited the activation of inflammatory leukocytes.

Here we report changes in expression patterns in intestinal tissue. The majority of research into RIC has focused on the heart and to a lesser extent the brain. Therefore the understanding of the processes within intestinal tissue which are important in the context of NEC, are not well understood and is the focus of our studies.

## Results

### RNA quality

The RIN scores for each sample are shown in Table 1: 23 out of the 24 samples had RIN scores greater than 8. The Concentration of RNA in each sample ranged from 1620 to 5060 ng/µl.

**Table 1:** Quality of RNA in each sample. Data provided by Qiagen. RIN >8 indicates high quality sample.^10,11^.

| Sample | Concentration of RNA (ng/μl) | RIN |
| --- | --- | --- |
| 1 | 1990 | 8.7 |
| 2 | 2460 | 9.4 |
| 3 | 2940 | 8.3 |
| 4 | 2540 | 9.2 |
| 5 | 3520 | 8.1 |
| 6 | 2920 | 9.1 |
| 7 | 2760 | 9.5 |
| 8 | 3780 | 9.0 |
| 9 | 2220 | 9.4 |
| 10 | 2760 | 9.5 |
| 11 | 3360 | 7.4 |
| 12 | 2780 | 9.3 |
| 13 | 2800 | 8.1 |
| 14 | 2980 | 9.5 |
| 15 | 2220 | 9.5 |
| 16 | 5060 | 8.2 |
| 17 | 4500 | 8.9 |
| 18 | 4540 | 8.2 |
| 19 | 3600 | 9.5 |
| 20 | 1620 | 9.3 |
| 21 | 2860 | 9.7 |
| 22 | 3900 | 9.1 |
| 23 | 2820 | 10.0 |
| 24 | 2800 | 8.9 |

### Data Exploration and Quality Control

Before proceeding to analysis of the differential gene expression, we excluded the lowly expressed mRNAs and then used hierarchical clustering, and PCA to ensure we had no outliers in the dataset indicating a technical issue with the RNA processing and the NGS. Figure 1 shows box plots of the raw sample counts of mRNA hits and the same dataset filtered to exclude lowly expressed mRNAs. This filtering reduces the noise from lowly expressed mRNA and thus the need for statistical correction of multiple testing. Hence this reduces the risk of a Type II statistical error. These charts also show very similar mRNA counts across all samples.

**Figure 1.**
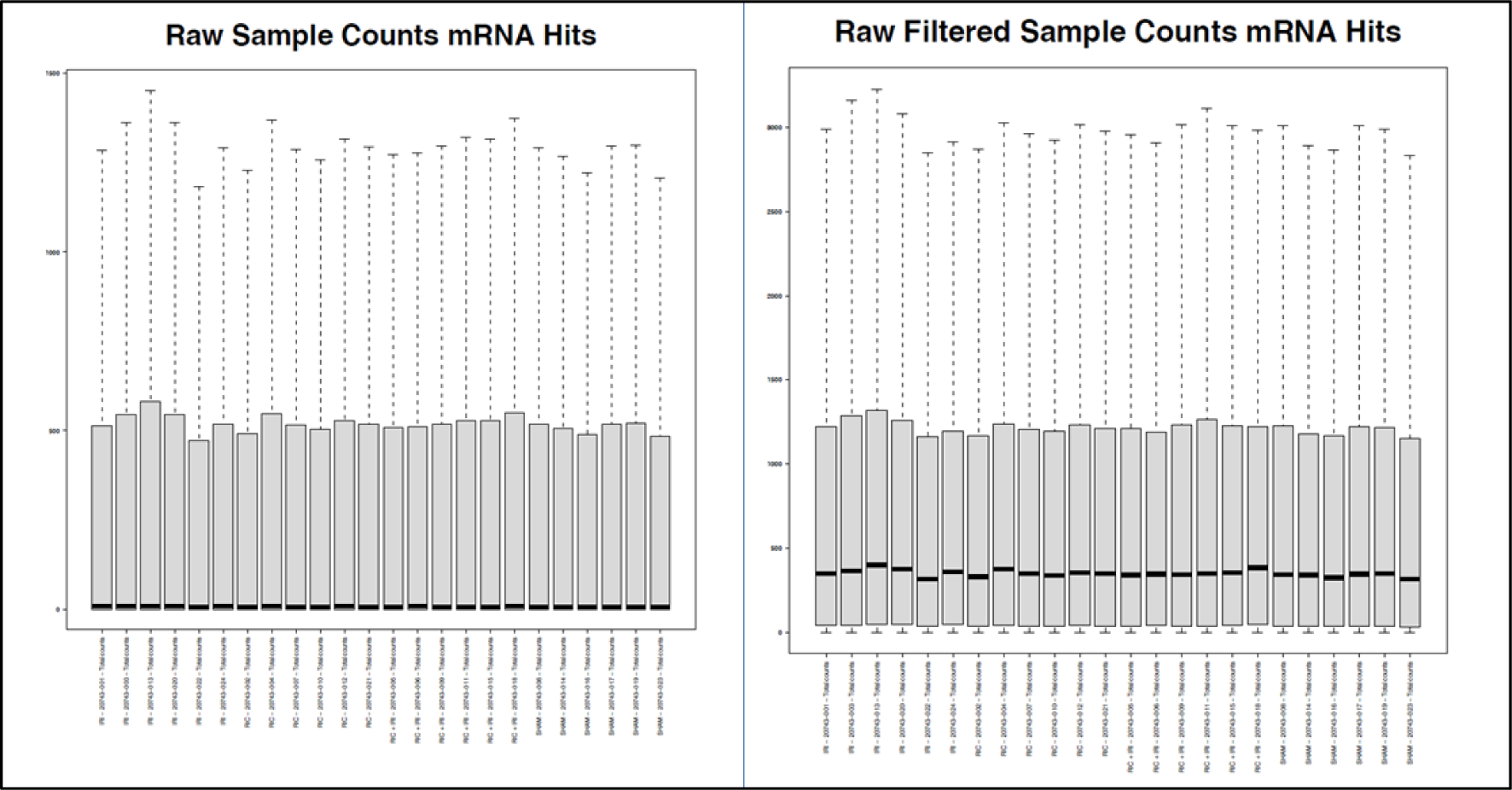
Box plot of Unfiltered and Filtered Sample Counts of mRNA Hits in each sample; The filter removes lowly expressed mRNAs. Median shown by black gene. Gene counts range from 0-1500 copies. Box shows 1.5x IQR. This is done to reduce noise and prevent Type II error.

The hierarchical clustering (using Euclidean distance and Average linkage) showed that in the whole dataset the IRI vs SHAM dominates over RIC vs SHAM in the patterns of gene expression, with samples from IRI and RIC+IRI clustering together and SHAM and RIC+SHAM clustering together (Figure 2). The PCA was labelled by known phenotypical factors (Age in days, gender, weight, litter, and date of procedure) there was no clustering by any of these specific factors. Sample 3 was a potential outlier in the PCA but not within raw counts/IQR plot. On the basis of these analyses, no samples needed to be excluded from further analysis of differential gene expression. (Figures 3 and 4).

**Figure 2.**
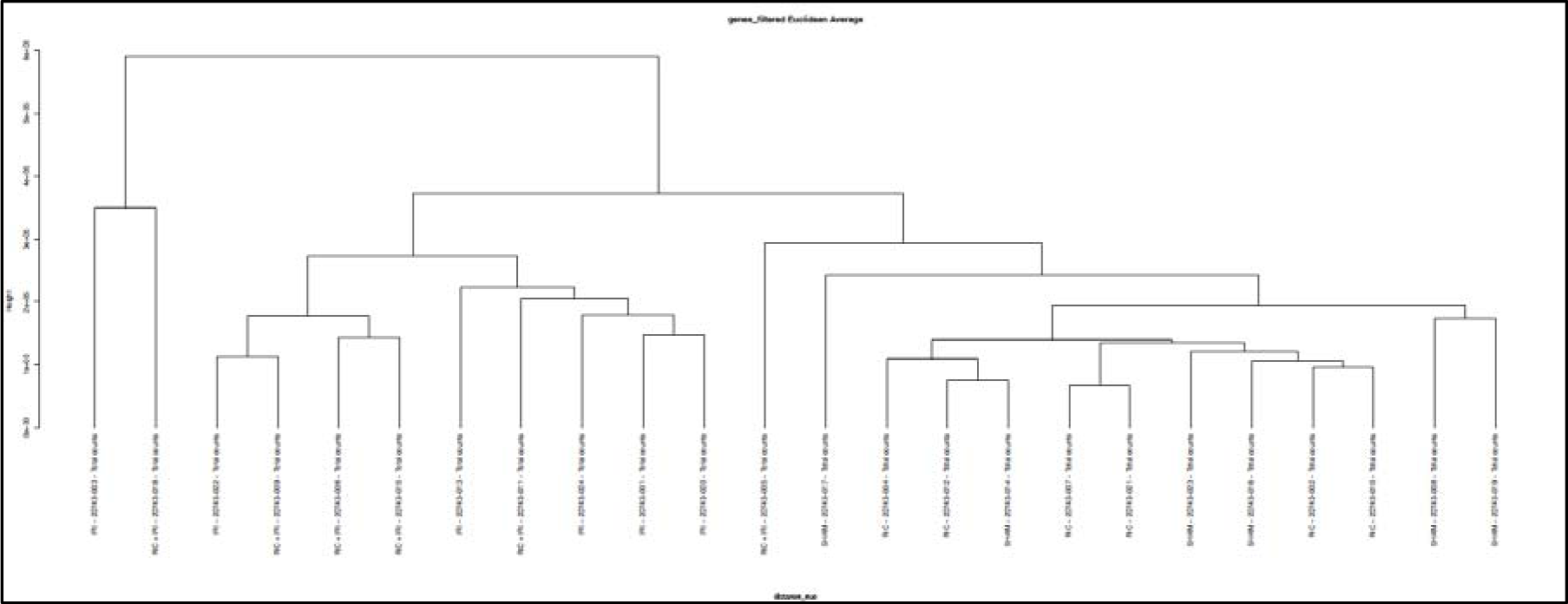
Hierarchical clustering of all samples from an RNA-Sequencing experiment which compared four conditions (list them). Using Euclidean distance and average linkage clustering showed that IRI vs No predominately defined the clusters in the whole model.

**Figure 3.**
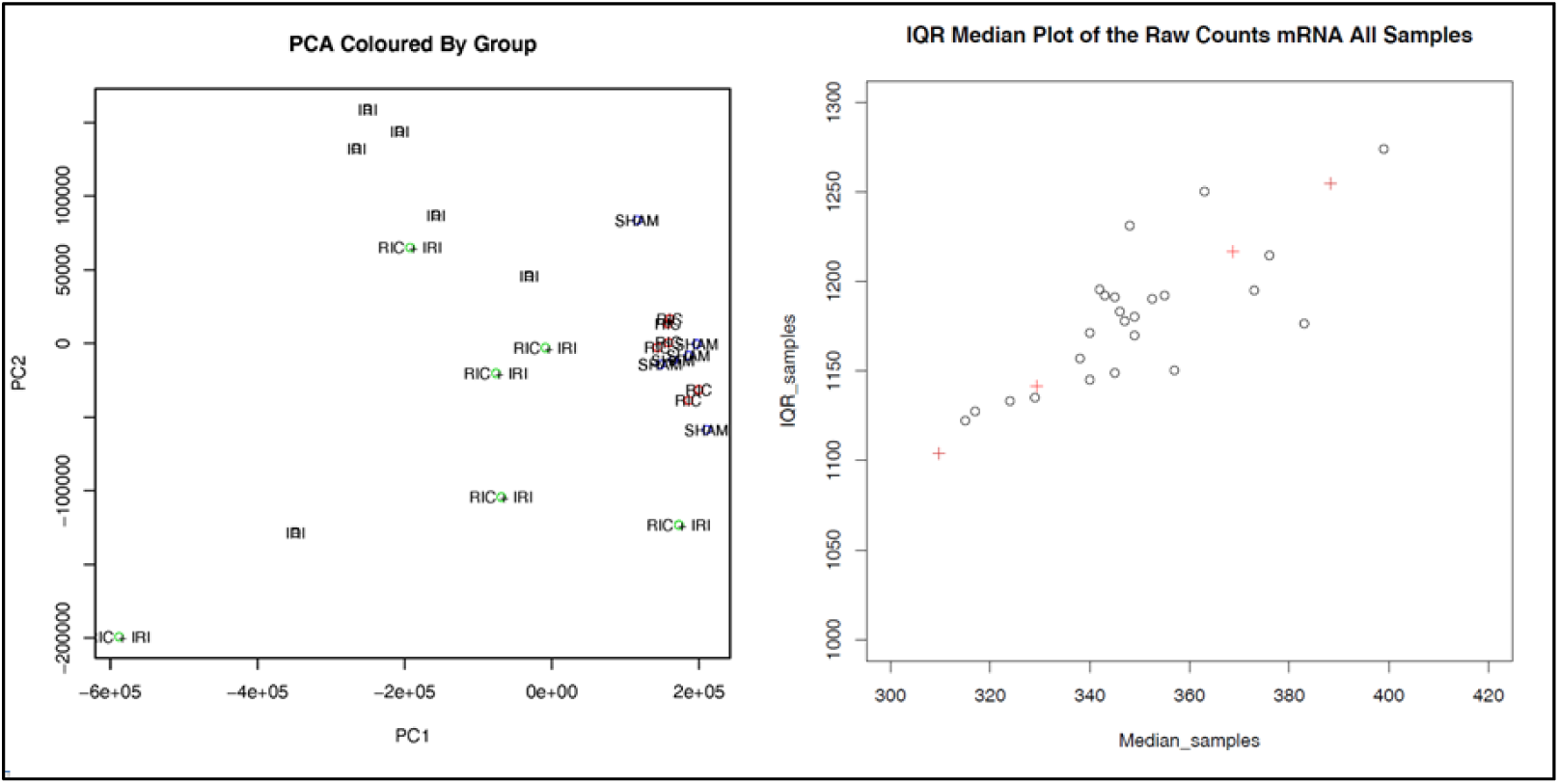
Principal component analysis (PCA) of all samples labelled by group. Median or Raw Counts plotted against Interquartile range of all samples and genes.

**Figure 4.**
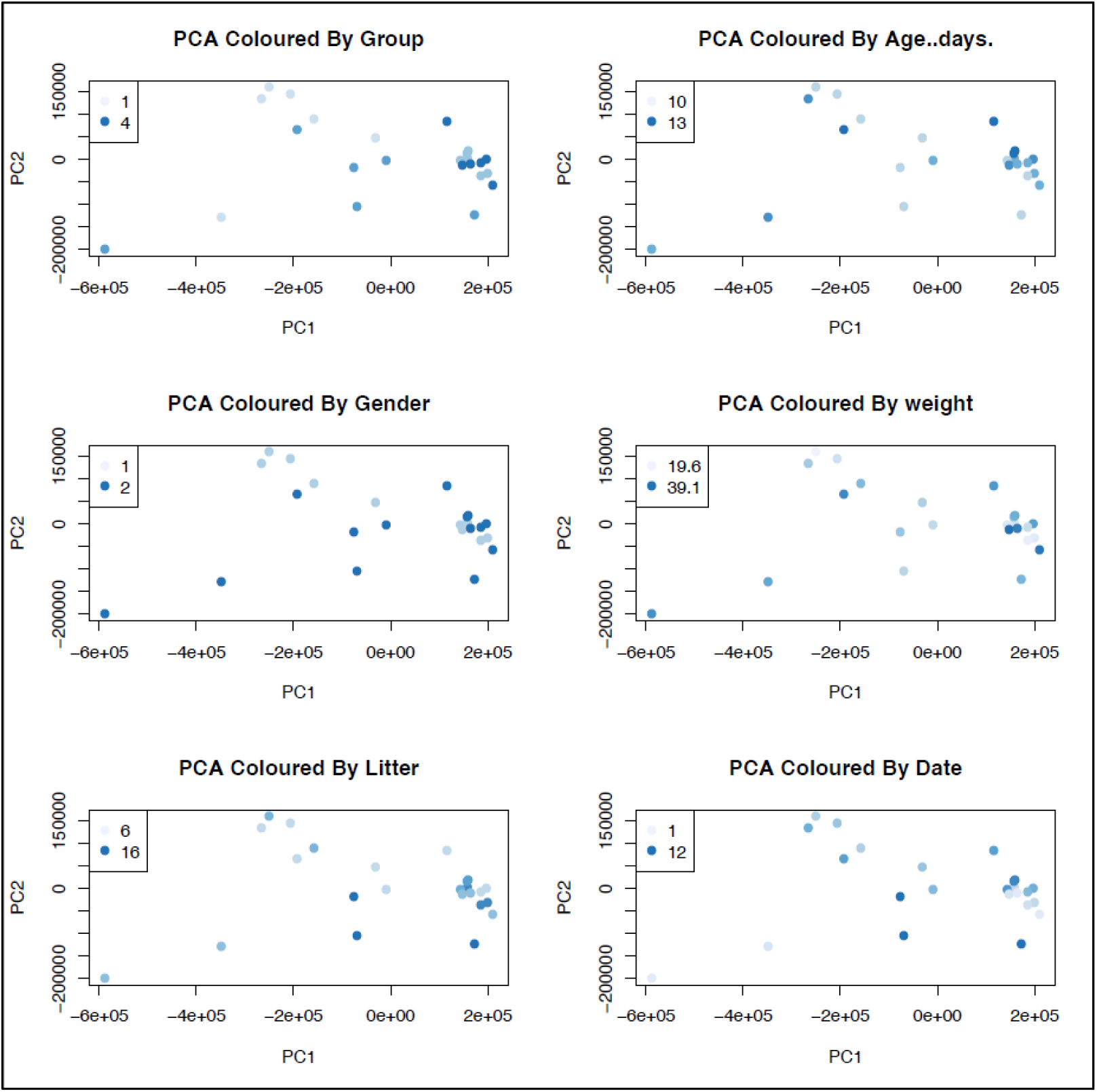
Principal component Analysis (PCA) of RNA Sequencing data of all 4 groups. These graphs showing that the known potential confounding factors are not showing a measurable effect. Factors examined: Age of pups; gender; weight; litter the pups were from; date of experiment.

### Differential Expression Testing

In order to address the key biological question of what is different in the intestinal tissue following conditioning that provides a protective effect. The most obvious analysis to carry out therefore is between animals that have undergone the IRI insult to the bowel (IRI only) and those that had conditioning prior to the injury (RIC+IRI). It is expected that this analysis will be informative but the tissue in question has undergone a significant physiological stress and damage and thus whether the RNA recovered gives meaningful data was not a certainty. Moreover, whilst the control group for the comparison has also undergone the same insult, this will inevitably have diverse and extensive effects on multiple biochemical pathways. The potential for a lot of noise here means that analysis of the effect of RIC without the IRI is also desirable. Conversely, this analysis alone may not be fully informative as it is at least conceivable that the protective processes (in terms of meaningful changes in gene expression) in the end organ are triggered both by the conditioning and the stimulus of the ischaemic insult. Hence the decision to focus on both a comparison between animals who underwent RIC vs controls and animals who underwent RIC prior to IRI vs IRI alone. This was done using edgeR.^11,12^ The model was built using the whole dataset and then the direct comparisons between the groups performed. *gplots* v 3.1.1,^17^ *edgeR*^18^ and *biomaRt*^19^ packages were used for these analysis. Volcano plots and heatmaps were produced to display the differentially expressed genes. biomaRt allows the gene IDs to be converted to gene and protein name.

We report three different two group comparisons: IRI vs SHAM; RIC+IRI+IRI; and RIC vs SHAM. The full tables of differentially expressed genes are included in the additional material. In animals that underwent IRI compared to SHAM 6772 genes showed statistically significant differential expression (q value <0.05) (Figure 5). In animals that underwent RIC and IRI (compared to IRI alone) there were 135 genes that showed differential expression (Figure 6) and the comparison between RIC and SHAM showed 868 genes with statistically significant gene expression (Figure 7). Figure 8 and table shows the overlap in genes that showed differential expression in both the RIC vs SHAM and RIC + IRI vs IRI groups.

**Figure 5.**
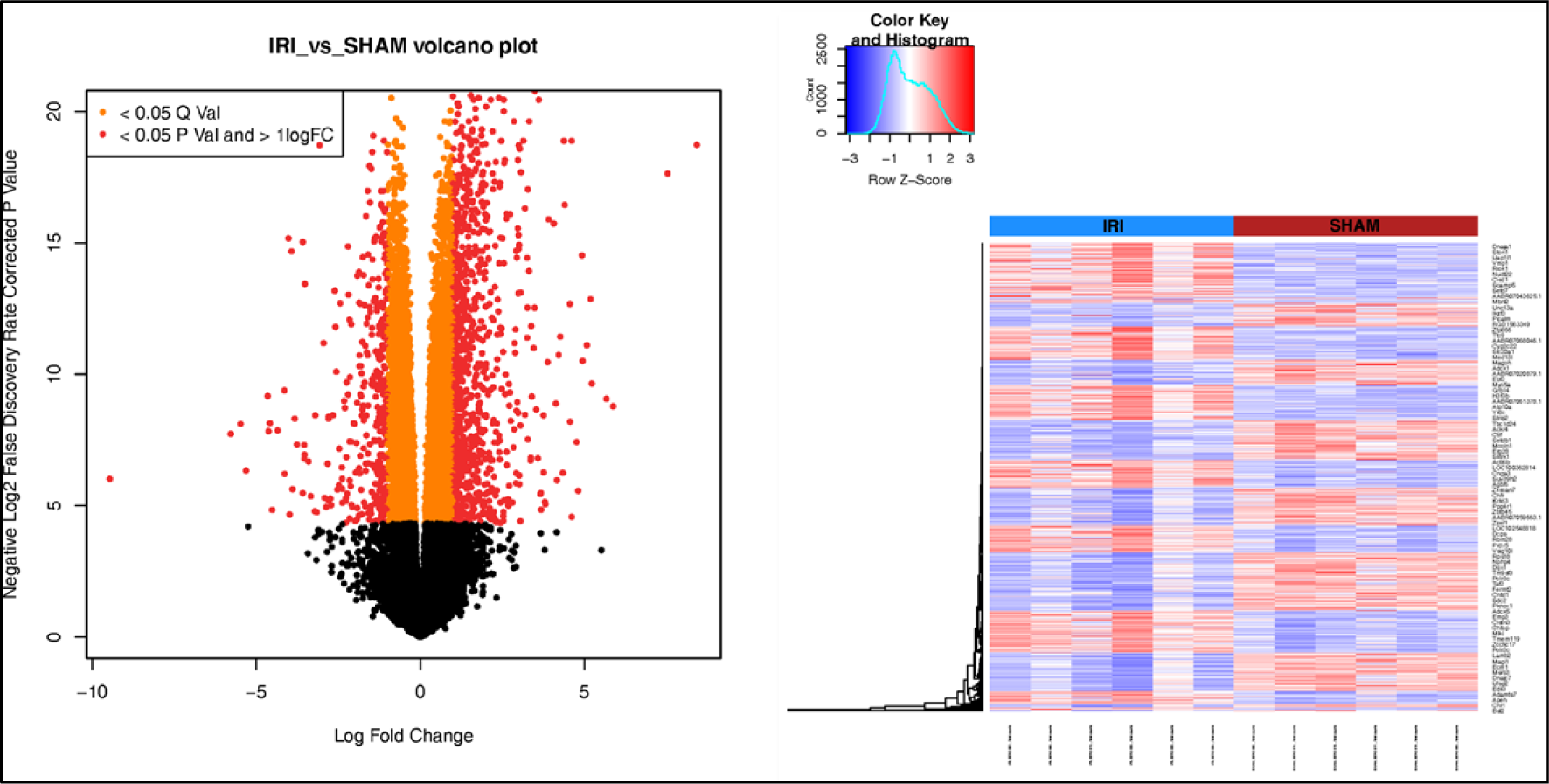
Vocano plot and Heat Map of IRI vs SHAM. Vocano plot shows LogFold Change in expression for all genes; orange colour indicates statistical significant (Q <0.05); red colour indicates a logFold change of greater than 1 and statistical significance. Heatmap shows all genes that showed signficance difference between groups: Blue indicates reduced expression; red increased. These data indicate the change in gene expression due to exposure to IRI alone.

**Figure 6.**
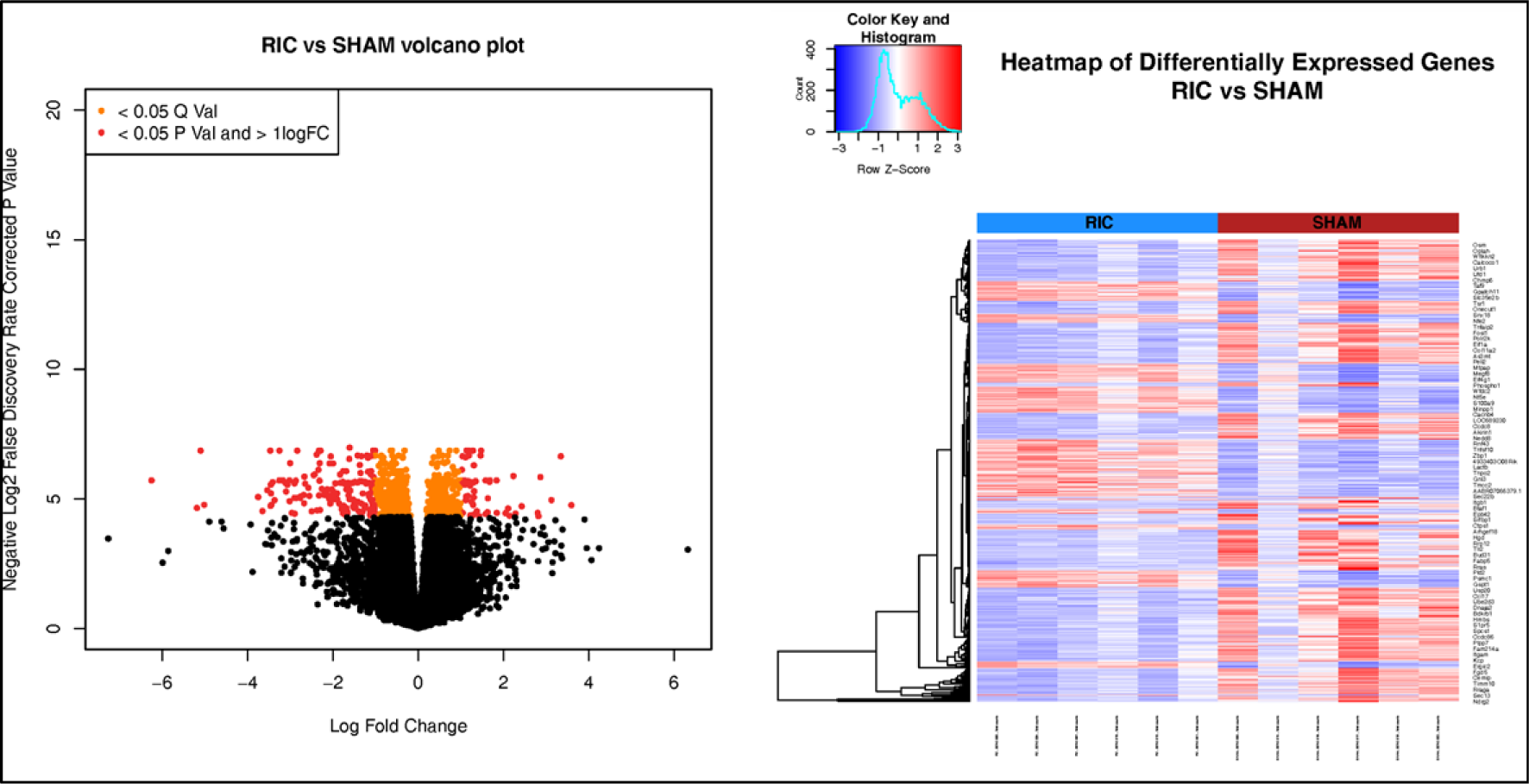
Vocano plot and Heat Map of RIC vs SHAM. Vocano plot shows LogFold Change in expression for all genes; orange colour indicates statistical significant (Q <0.05); red colour indicates a logFold change of greater than 1 and statistical significance. Heatmap shows all genes that showed signficance difference between groups: Blue indicates reduced expression; red increased. These data indicate the change in gene expression due to exposure to RIC alone.

**Figure 7.**
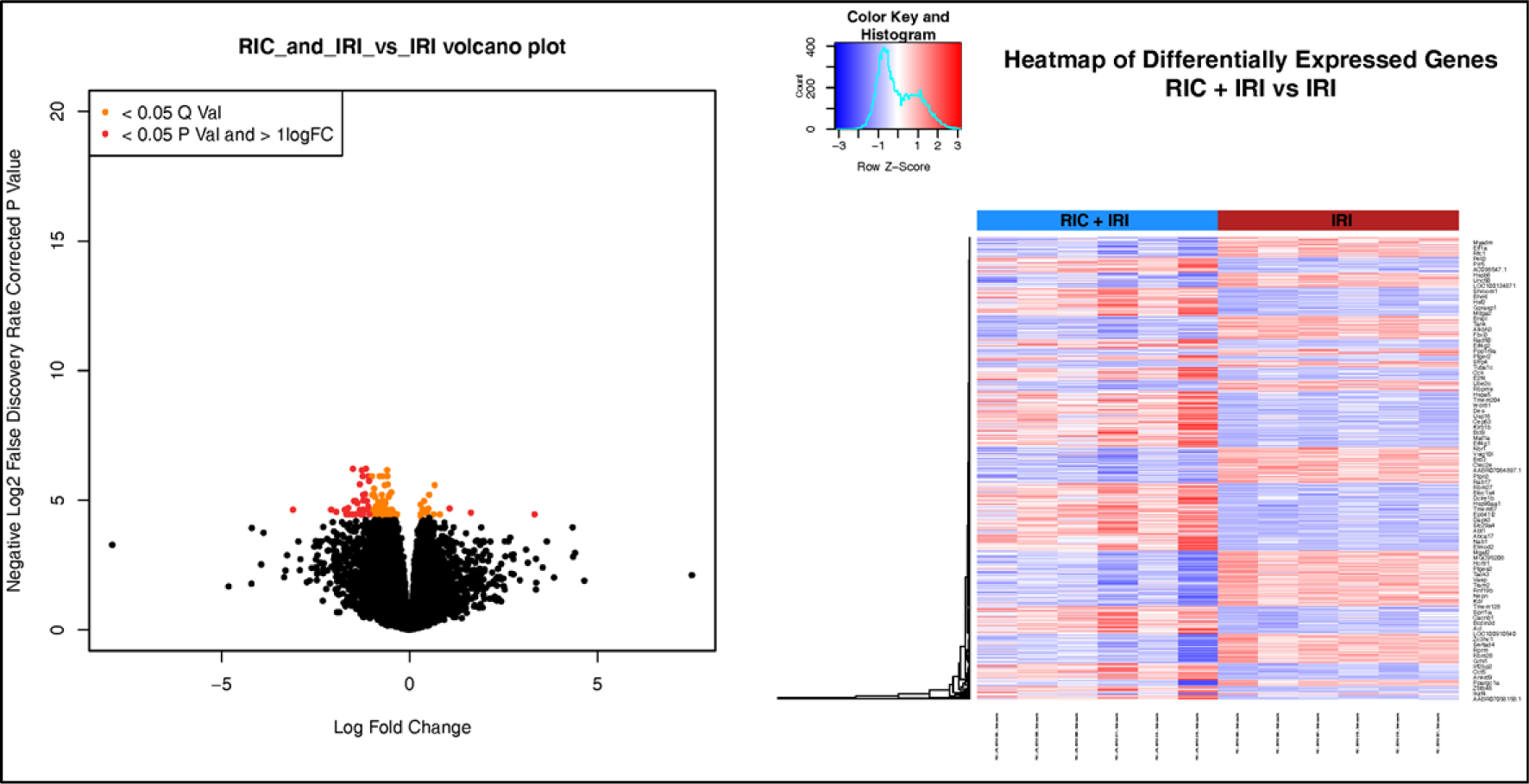
Vocano plot and Heat Map of RIC+IRI vs IRI. Vocano plot shows LogFold Change in expression for all genes; orange colour indicates statistical significant (Q <0.05); red colour indicates a logFold change of greater than 1 and statistical significance. Heatmap shows all genes that showed signficance difference between groups: Blue indicates reduced expression; red increased. These data indicate the change in gene expression due to exposure to RIC in anaimals that also underwent IRI.

**Figure 8.**
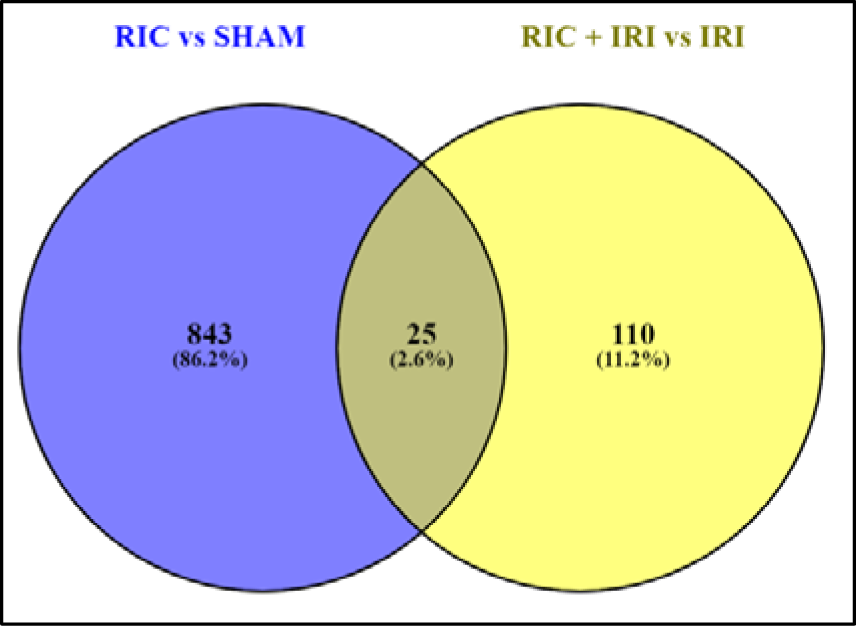
Venn diagram of genes showing statisically significant changes in expression two, two-way comparisons: RIC vs SHAM and RIC+IRI vs IRI. 25 genes showed changes in expression in both comparisons.

Among these 25 genes found in both bilateral comparisons (Table 2), are several genes which are likely candidates for involvement in a putative RIC pathway within the intestine as they are known to be important in inflammatory pathways.^23^ These include CXCL1 which is an important chemoattractant, TIMP1 which is anti-apoptosis, CD-55 which is known to be a complement regulator, SOD2 which is involved in the detoxification of reactive oxygen species and Nfκβ2 which is a key part of the innate inflammation pathway.^23^

**Table 2.** Short names of genes found to be differentially expressed in both RIC vs SHAM and RIC+IRI vs IRI.

|  |  |  |  |  |  |
| --- | --- | --- | --- | --- | --- |
| 1 | Lrrc8c | 10 | Malsu1 | 18 | Map3k8 |
| 2 | Ugdh | 11 | Spry2 | 19 | Sod2 |
| 3 | Cxcl1 | 12 | Timp1 | 20 | Nfkb2 |
| 4 | Adamts4 | 13 | Tfpi2 | 21 | Procr |
| 5 | Cd55 | 14 | Plau | 22 | Usp36 |
| 6 | Trib1 | 15 | Coq10b | 23 | Kcne4 |
| 7 | Adamts8 | 16 | S1pr3 | 24 | Gstm5 |
| 8 | Pde4b | 17 | Ifitm3 | 25 | AABR07047256.1 |
| 9 | RGD1563365 |  |  |  |  |

### Differential Expression – Targeted Gene Analysis

Tables 3-5 show the results of the targeted gene analysis. In terms of cell markers, RIC significantly reduced the expression of Macrophage markers (Figure 11). CD3 was reduced by RIC suggesting a reduction in T-cells but both CD4 and CD8a were not statistically significantly changed (Figure 12). Both Fibroblast and alpha-smooth muscle markers were increased by IRI and this effect was mitigated by RIC (Figure 13).

**Table 3.** Four way comparison of gene expression of specific markers: Cell type markers.

| Cell Type | Marker Used | Result | p-value |
| --- | --- | --- | --- |
| Immune Cells | CD45 | No Change |  |
| Macrophages | CD163 | Increased by RIC | 0.003 / <0.001 |
|  | CD164 | No Change |  |
|  | CD206 | Reduced by RIC AND IRI combined | 0.006 |
|  | CD68 | No Change |  |
|  | CD11C | Reduced by RIC | 0.014 / <0.001 |
| B-cells | CD20 | No Change |  |
|  | CD19 | No Change |  |
| T-cells | CD3 | Reduced by RIC | 0.019 / 0.047 |
|  | CD4 | No Change |  |
|  | CD8a | No Change |  |
|  | CD69 | Increased by IRI | 0.002 / 0.003 |
|  | CD103 | No Change |  |
| Natural Killer Cells | CD56 | Increased by RIC, Increased by IRI | < 0.001 / 0.029 |
|  | KIR | No Change |  |
| Endothelial cells | CD109 | No Change |  |
| Neutrophils | CD15 | Reduced by IRI (Effect mitigated by RIC) | <0.001 / 0.043 / 0.003 |
| Fibroblasts | VIM | Increased by IRI (Effect mitigated by RIC) | <0.001 / 0.022 / 0.025 |
|  | CDH2 | No Change |  |
|  | ACTA 2 | Increased by IRI, Reduced by RIC | 0.001 / 0.02 / 0.043 |
| Alpha Smooth Muscle | PAFAH1B3 | Increased by IRI (Effect mitigated by RIC) | 0.049 |
|  | DESMIN | Increased by IRI, Reduced by RIC | 0.01 / <0.001 / 0.013 / 0.013 |
|  | CDH2 | No Change |  |

**Table 4.** Four way comparison of gene expression of specific markers: Hypoxia, cell toxicity and cell repair.

| Biological Processes | Marker Used | Result | p-value |
| --- | --- | --- | --- |
| Hypoxia | HIF-1a | Increased by IRI, Reduced by RIC | <0.001 / 0.005 / <0.001 |
|  | HIF-1b | No Change |  |
|  | HIF-2a | Increased by IRI, Reduced by RIC | 0.038 / 0.009 / 0.023 / <0.001 |
|  | VEGFa | Increased by IRI (Effect mitigated by RIC) | <0.001 / 0.041 / <0.001 |
| Cell Toxicity | GZMA | No Change |  |
|  | GZMB | No Change |  |
| Cell Repair | KRT-7 | Increased by IRI (Effect mitigated by RIC) | 0.006 / 0.049 / 0.01 |
|  | KRT-8 | Increased by IRI | 0.012 / 0.02 |
|  | KRT-18 | Increased by IRI (Effect mitigated by RIC) | 0.001 / 0.003 / 0.004 |
|  | KRT-19 | Increased by IRI (Effect mitigated by RIC) | 0.018 / 0.016 |

**Table 5.** Four way comparison of gene expression of specific markers. Specific biological pathways; Molecular pathways known to be involved in NEC pathophysiology and molecules that form the RISK pathway in cardiac iscaehemic conditioning.

| Specific Biological Pathways | Marker Used | Result | p-value |
| --- | --- | --- | --- |
| Molecules known to be involved in NEC pathophysiology | TLR2 | Increased by IRI, Reduced by RIC | 0.042 / 0.044 / 0.005 / 0.006 |
|  | TLR4 | Increased by IRI | 0.005 / 0.005 |
|  | MyD88 | Increased by IRI | 0.002 / 0.001 |
|  | NF-KB1 | Increased by IRI, Reduced by RIC | 0.032 / 0.006 / 0.019 / 0.003 |
|  | CD17 | No Change |  |
|  | iFABP | No Change |  |
| Molecules known to be involved in RIC in cardiac tissue | MEK1 | Reduced by IRI | 0.002 / 0.032 |
|  | MEK2 | Reduced by IRI, Reduced by RIC | 0.009 / <0.001 |
|  | P38 | Reduced by IRI | 0.001 / 0.012 |
|  | P13K | Increased by RIC | 0.023 / <0.001 |
|  | PKC | Reduced by IRI | <0.001 / 0.01 |

HIF1-α, HIF2-α and VEGFα – markers of hypoxia – were all statistically significantly increased by IRI and reduced by RIC (Figure 14). Similarly, Markers of cell-repair mechanisms (KRT-7,-8,-18 and -19) were increased in animals exposed to IRI with this effect mitigated by RIC (Figure 15).

Toll-like Receptor-2, Toll like receptor-4, MyD88 and Nfκβ1 were all increased by IRI. This effect was mitigated by RIC (Figure 16). MEK1, MEK2, P48 and PKC were all reduced by IRI, suggesting a role in ischaemic injury. PI3K in increased with RIC. (Figure 17)

## Conclusions

The protective effect of RIC is well-established in multiple species and multiple organ systems.^3^ In this work we showed a profound reduction in intestinal injury in rat pups exposed to IRI.^4^ The mechanisms by which RIC confers this protective effect are complex and significant work over the past decades has elucidated multiple potential pathways involved. Within the context of the intestine, the ‘end-organ’ pathway has not been well studied. Utilising transcriptomics, we sought to better understand how RIC confers such a profound reduction in intestinal injury.

### Differential Expression testing

Differential expression was seen in 868 genes in the RIC vs SHAM group and 135 in the RIC+IRI vs IRI alone comparison. Genes that are differentially expressed in both groups are likely to be important. There are 25 genes that are differentially expressed in both groups (Table 2). Of these 25 genes, eight are known to have to be involved with functions likely to be important in the mechanisms of RIC. Table 6 shows these eight genes and the results from the differential expression analysis; specifically the fold change in the expression levels and the corrected p-values.

**Table 6.** Genes showing differential expression in both the RIC vs SHAM and RIC+IRI vs IRI groups. *logFC = log fold-change in gene expression; logCPM = log Counts per Million; PValue = raw p-value; FDR = False discovery rate (p-value corrected for multiple comparisons)*.

| Genes |  | logFC | logCPM | PValue | FDR |
| --- | --- | --- | --- | --- | --- |
| **CXCL1** | *RIC+IRI* | -0.73581 | 8.295629 | 9.37E-06 | 0.016462 |
|  | *RIC* | -0.59385 | 7.581155 | 6.20E-05 | 0.014521 |
| **TIMP1** | *RIC+IRI* | -1.08748 | 4.695321 | 9.35E-05 | 0.036933 |
|  | *RIC* | 0.584135 | 6.188187 | 0.00031 | 0.021415 |
| **Cd55** | *RIC+IRI* | -0.56804 | 7.283793 | 1.78E-05 | 0.020411 |
|  | *RIC* | 0.487938 | 6.174565 | 9.45E-05 | 0.016591 |
| **Map3k8** | *RIC+IRI* | -0.75824 | 4.211773 | 0.000186 | 0.042433 |
|  | *RIC* | -2.7615 | 2.360486 | 0.000722 | 0.030387 |
| **Sod2** | *RIC+IRI* | -0.98012 | 7.372327 | 0.000217 | 0.043591 |
|  | *RIC* | -1.47528 | 3.472257 | 0.000946 | 0.033794 |
| **Nfkb2** | *RIC+IRI* | -0.49015 | 4.366177 | 0.000222 | 0.043961 |
|  | *RIC* | -0.3912 | 4.601592 | 0.000969 | 0.034104 |
| **Procr** | *RIC+IRI* | -1.34504 | 0.765029 | 0.000227 | 0.044383 |
|  | *RIC* | -0.40365 | 7.534098 | 0.000969 | 0.034104 |
| **Usp36** | *RIC+IRI* | -1.60342 | -1.88456 | 0.000267 | 0.045566 |
|  | *RIC* | -0.80179 | 2.778097 | 0.001428 | 0.040053 |

**Table 7.**
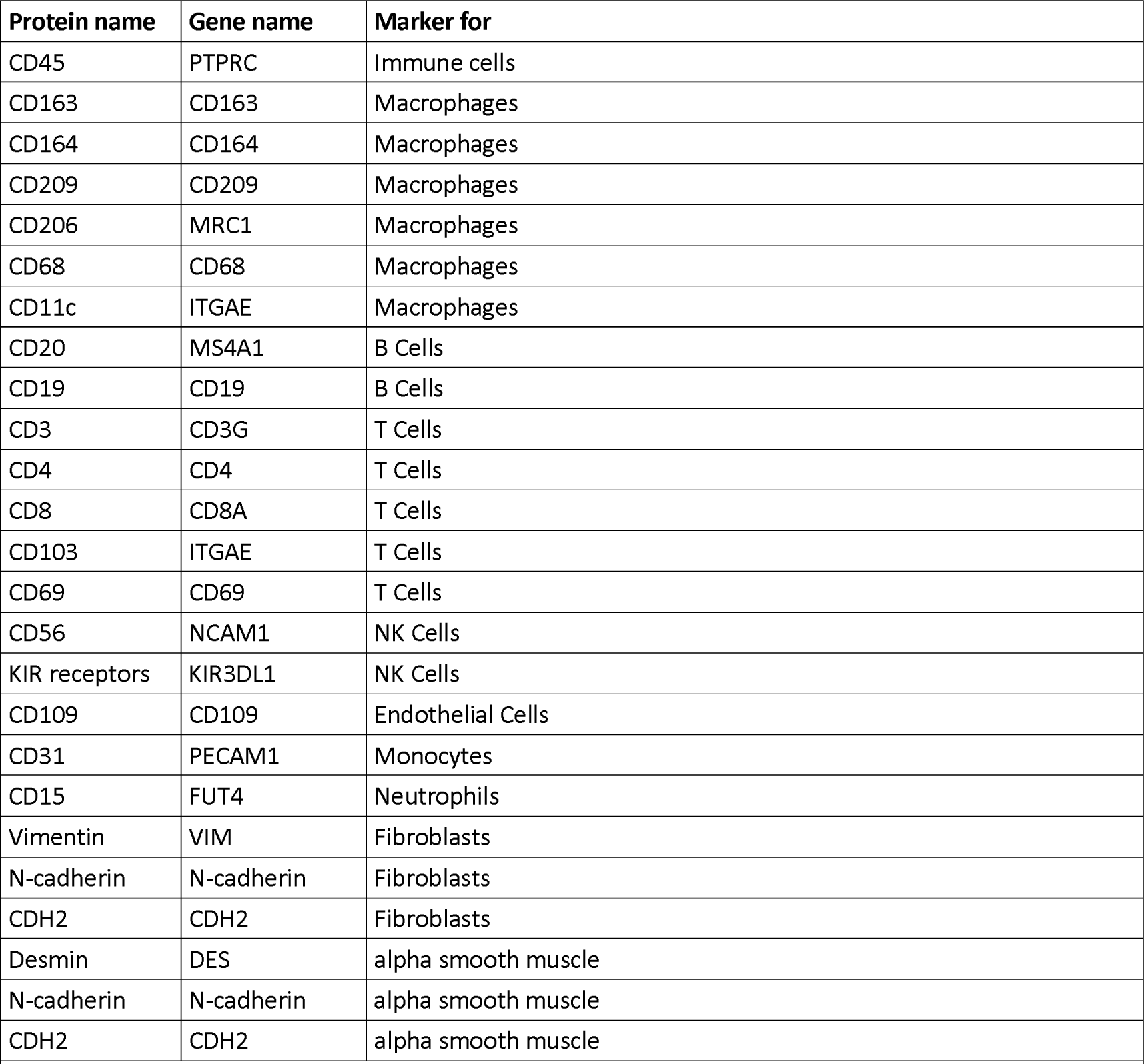
Selected proteins used as markers of the cell populations. *The gene names refer to the names used by the Ensembl database.^20^*

| Protein name | Gene name | Marker for |
| --- | --- | --- |
| CD45 | PTPRC | Immune cells |
| CD163 | CD163 | Macrophages |
| CD164 | CD164 | Macrophages |
| CD209 | CD209 | Macrophages |
| CD206 | MRC1 | Macrophages |
| CD68 | CD68 | Macrophages |
| CD11c | ITGAE | Macrophages |
| CD20 | MS4A1 | B Cells |
| CD19 | CD19 | B Cells |
| CD3 | CD3G | T Cells |
| CD4 | CD4 | T Cells |
| CD8 | CD8A | T Cells |
| CD103 | ITGAE | T Cells |
| CD69 | CD69 | T Cells |
| CD56 | NCAM1 | NK Cells |
| KIR receptors | KIR3DL1 | NK Cells |
| CD109 | CD109 | Endothelial Cells |
| CD31 | PECAM1 | Monocytes |
| CD15 | FUT4 | Neutrophils |
| Vimentin | VIM | Fibroblasts |
| N-cadherin | N-cadherin | Fibroblasts |
| CDH2 | CDH2 | Fibroblasts |
| Desmin | DES | alpha smooth muscle |
| N-cadherin | N-cadherin | alpha smooth muscle |
| CDH2 | CDH2 | alpha smooth muscle |

**Table 8.** Selected proteins used as markers of hypoxia and cellular injury. *The gene names refer to the names used by the Ensembl database.^20^*

| Protein name | Gene name | Marker for |
| --- | --- | --- |
| HIF1a | HIF1a | Hypoxia |
| HIF2a | ARNT | Hypoxia |
| HIF1b | EPAS1 | Hypoxia |
| VEGF | VEGFA | Hypoxia |
| KRT7 | KRT7 | Type 1 cytokeratins – cell repair |
| KRT8 | KRT8 | Type 1 cytokeratins – cell repair |
| KRT18 | KRT18 | Type 2 cytokeratins – cell repair |
| KRT19 | KRT19 | Type 2 cytokeratins – cell repair |

**Table 9.**
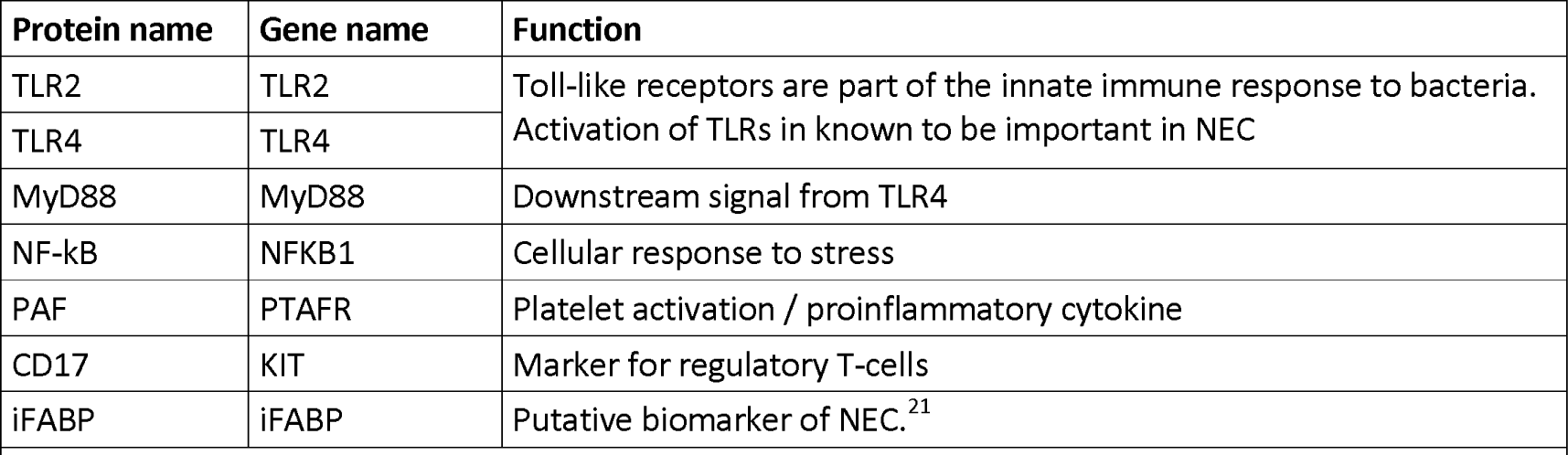
Selected proteins shown to be important in the pathophysiology of NEC. The gene names refer to the names used by the Ensembl database.^20^

**Table 10.**
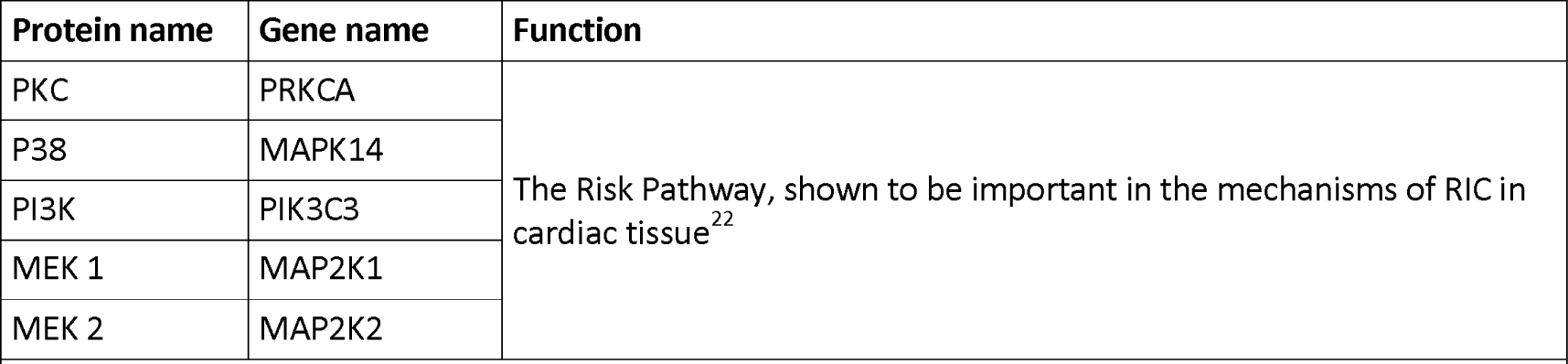
Selected proteins shown to be important in the mechanisms of RIC in other target organs. The gene names refer to the names used by the Ensembl database.^20^

C-X-C Motif Chemokine ligand-1 (Cxcl-1) is a neutrophil chemoattractant^23^ that is downregulated by RIC. A Pubmed search showed no previous reports of a role of Cxcl-1 in the mechanisms of RIC. Interestingly, it has been established as an important part of the pathophysiology of brain damage after stroke. The production of Cxcl-1, through the NF-ĸβ-1 dependent pathway triggers neutrophil infiltration and thus leads to neuroinflammation.^24^ Similarly, Cxcl-1 has been shown to recruit neutrophils in cardiac ischaemia.^25^ Reduced expression of Cxcl-1 due to RIC, leading to less neutrophil recruitment in the presence of IRI, leading to less inflammation is a very plausible mechanism. Thus the effect of RIC maybe to reduce neutrophil mediated tissue damage.

Mitogen-Activated Protein Kinase Kinase Kinase 8 (MAP3k8), as with many of the genes discussed here, has an important role in oncogenesis. Map3k8 also has some interesting downstream effects. It induces the production of NF-ĸβ as well as the production of TNF-α and IL-2.^23^ NF-ĸβ is discussed below but this shows consistent changes in expression at three different stages in the pathway; MAP3k8 induces NF-ĸβ and one of the downstream effects of NF-ĸβ is to increases the expression of Cxcl-1. In theses data the expression of all three is suppressed.

Nuclear factor kappa-light-chain-enhancer of activated B cells pathway subunit 2 (NF-ĸβ2) is part of the NFĸβ family of transcription factors composed of five structurally related members including NF-ĸβ1 and NF-ĸβ2. ^28^ Functional NF-ĸβ2 is produced by the post-translational processing of its precursor protein p100. This may be especially important in the so-called ‘non-canonical’ NF-ĸβ pathway.^29^ The precursor protein (p100) has an inhibitory function on NF-ĸβ^28^ and thus changes in the RNA expression levels could have promoter or inhibitory effects on the whole pathway, depending on whether the p100 protein is rapidly converted to mature NF-ĸβ2 or not. However, the Map3k8 and Cxcl-1 expression levels along with NF-ĸβ2 are all reduced in RIC suggesting that the overall effect is to inhibit NF-ĸβ pathway.

Superoxide Demutase 2 (SOD2) is an important part of the cells response to oxidative stress and dismutates superoxide to hydrogen peroxide.^26^ Hence, the reduced expression of SOD2 by RIC is at first glance surprising. However, increased production of hydrogen peroxide through the up-regulation of SOD2 stimulates pro-oxidants involved in apoptosis. Thus reduced expression of SOD2 can be said to be anti-apoptotic.^27^

Protein C Receptor (Procr)is a receptor for activated Protein C known as endothelial cell protein C receptor (EPCR).^23^ Activated Protein C is an important inhibitor of the clotting cascade but it is also exerts important anti-inflammatory effects.^30^ The Protein C Receptor protein is found on the surface of endothelial and other cells types. Down-regulation of this receptor (seen in both RIC and RIC and IRI groups) implies a dampening down of Activated-Protein C suppression of coagulation. Coagulation is a cardinal feature of ischaemic necrosis.^31^ Activated-protein C, via the Protein C receptor, triggers anti-inflammatory downstream effects via several mechanisms including suppressing NF-ĸβ. ^30,32^ Therefore, the down-regulation of this receptor in these data is potentially surprising, however it suggests other mechanisms are capable of reducing NF-ĸβ activation in this complex system, so lowering EPRC may be beneficial in this context. In addition, thrombin activation or EPCR produces a pro-inflammatory response and a disruption of the endothelial barrier.^32,33^ Hence, EPCR has a pro-inflammtory mechanisms quite apart from the binding ligand that it is named for and reduced expression of this molecule would be expected to result in reduced inflammation. Although the cross-talk for this multi-ligand receptor is undoubtedly complex.^29^

Ubiquitin Specific Protease 36 (Usp36) could be a putative part of the RIC protective pathway because of its suggested role in autophagy.^23^ Autophagy, a mechanism by which cells removes unnecessary or damaged components is thought to be important in maintaining cellular function in response to stress.^34^ Loss of Usp36 function autonomously activates autophagy.^35^ This implies that reduced expression of Usp36 would cause the cell to increase its autophagic activity which could be important in responding to the severe stress of IRI. Conversely, work in the mouse and human kidney suggests that Usp36 interacts with SOD2 in the mechanism of acute kidney injury due to ischaemia. These data suggests that increased Usp36 expression would be protective.^36^ appears to have multiple functions. One such function is that it may be anti-apoptotic.^23^ In these data, its expression is increased in RIC (compared to SHAM) but decreased in animals that undergo RIC and IRI. In the brains of rats, TIMP1 has been shown to be increased in response to infarction.^37^ Thus the decreased expression in animals exposed to RIC and IRI compared to IRI alone may be a reflection of the reduced injury. However, the increase seen in RIC alone is less easily explained. As the name implies, TIMP proteins are named for their inhibition of metalloprotinases (MMPs) which in turn play a key role in the normal physiology and wound healing of connective tissue.^38^ This inhibitory relationship is anti-inflammatory.

CD55 is a regulator molecule of the complement cascade. ^23^ Several studies looking at renal IRI have shown that over-expression of CD55 is protective against inflammation.^39^ Although a specific role with respect to RIC has not been established. As with TIMP1, these results show increased expression of CD55 in the presence of RIC but in animals that underwent IRI and RIC, the expression is reduced compared to IRI alone. Thus if CD55 is part of the protective effect of RIC (and it clearly has an anti-inflammatory role in other contexts), it modulation of inflammation (presumably via the complement cascade) is not straight-forward.

Nuclear factor kappa-light-chain-enhancer of activated B cells pathway (NF-ĸβ), discovered in 1986,^40^ has been extensively studied as a master regulator of pro-inflammatory genes,^28^ changes in the levels of TNF-α, IL-6 and IL-8 equivalent (KC/GRO) are seen in these animals. CXCL-1 showed reduced expression in both groups, indicating a concomitant down-stream effect of NF-ĸβ. Further corroboration is needed to confirm this but these data implicate a key role for NF-ĸβ in the protective effect of RIC in the intestine.

Toll-like Receptor-4 (TLR-4) triggers the NF-ĸβ, pathway.^41^ Importantly, this is an early step in the pathogenesis of NEC, prior to the development of IRI. Indeed proponents of an immunological understanding of NEC, often cite the role of TLR-4 as being central to the development of the disease.^42,43^ Ultimately, IRI is the common end-point of multiple pathways in NEC (although, it also leads to a vicious cycle of further inflammation and bowel compromise) thus one might expect RIC to only be effective at this late stage in the pathogenesis. Clearly, if RIC is effective in the intestine, it would be effective at reducing the ischaemic injury but if RIC is acting on NF-ĸβ as these data would suggest, then it is also likely to offer a protective effect at every stage in the pathogenesis of NEC. Whilst this hypothesis is not formally proven, it does present the prospect that RIC will be protective at multiple stages in the pathogenesis of NEC and thus (assuming that RIC can be delivered safely to human infants) could be a very effective therapy.

### Differential Expression – Targeting Gene analysis of cellular markers

Five cellular markers were used for macrophages: CD163, CD206 and CD11c showed changes that were statistically significant, whilst the others (CD163 and CD68) showed no change. CD163 was increased by RIC, whilst CD206 and CD11c were reduced (Figure 9). These different results suggest potentially that the cell population is not changing, rather the function of the macrophages is being altered. The regulation of these two proteins is different: CD163 is increased by IL-10, whilst CD206 is upregulated by IL-4 and IL-13^44^

**Figure 9.**
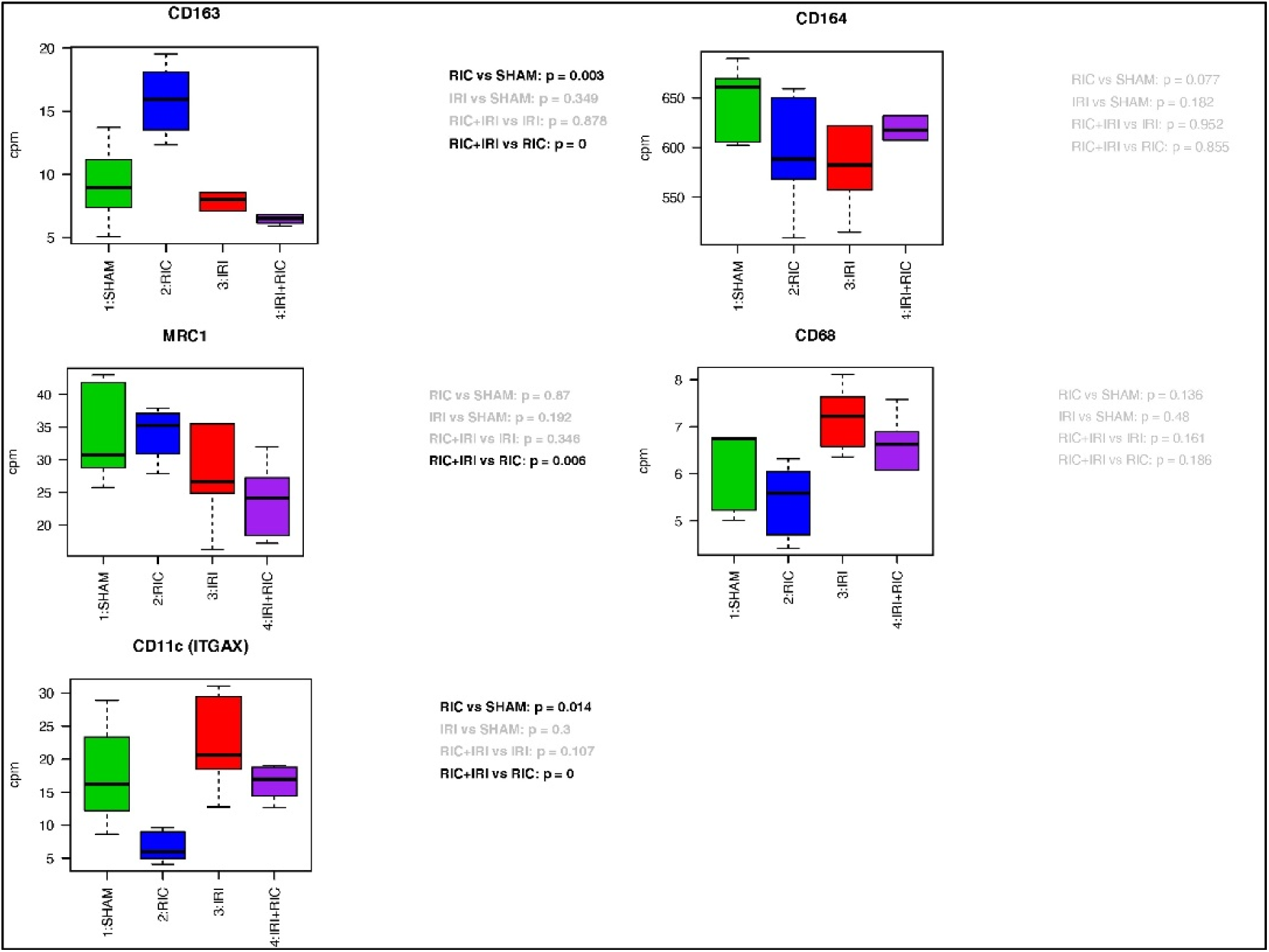
Four way comparison of gene expression of specific markers: Cell type markers for macrophages.

In addition, T-cell markers show an interesting pattern (Figure 10). CD4 and CD8a were unchanged. CD3g was reduced by RIC. There was a non-significant trend suggesting an increase with IRI. Animals exposed to IRI showed a reduction compared to IRI alone. One was to understand this is that there is an increase in T-cells due to IRI which is mitigated / prevented by RIC. Given that the CD4 and CD8a markers are unchanged this change would have to be due to neither T-Helper or cytotoxic T cells. The pattern of CD69 expression shows a statistical increase in animals exposed to IRI with non-significant trends from RIC. Taking these two results together would suggest that it might be regulatory T-cells that are important here. CD69 is also expressed by NK-cells but the other NK-cell markers used CD56 showed the opposite effect with RIC increasing the expression and KiR showing no change.

**Figure 10.**
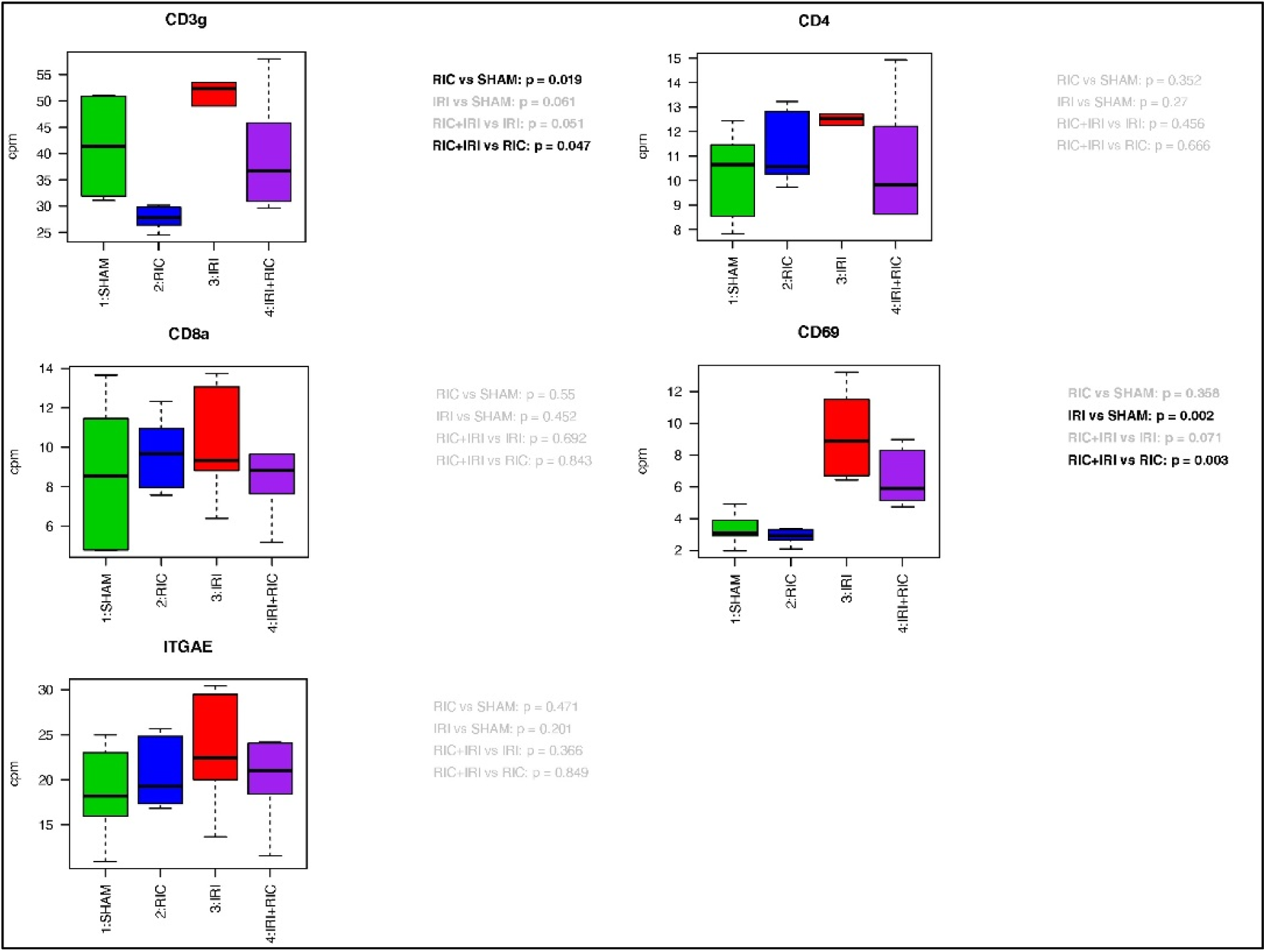
Four way comparison of gene expression of specific markers: Cell type markers for T-cells.

Biological processes involving the role of HIF-1 in ischaemia is well-established. HIF-1 is a transcription factor made of two subunits (HIF-1α and HIF-1β) and is a key part of the cell’s response to hypoxia. Hence it is a marker of hypoxic insult at the cellular level. Under normoxic conditions HIF-1α has very high turnover, being constitutively expressed and broken down by proteasomal degradation. Hypoxia inhibits the breakdown of HIF-1α thus leading to its nuclear accumulation.^45^ It has been shown that HIF-1α is increased in the intestinal injury of NEC,^46^ which is part of the evidence base supporting the notion that NEC is an ischaemia-reperfusion disease. In these animal experiments we demonstrated that HIF-1α expression seems to be following the pattern of the intestinal injury and this is mitigated by RIC: HIF-1α is increased with IRI and this effect it mitigated with RIC. HIF-1β is unchanged, whist HIF2α and VEGFα follow the same pattern as HIF-1α (Figure 12). Given the way that HIF-1 functions with constitutional expression of HIF-1β, it is logical that there is a change in the expression of HIF-1 α and not HIF-1β. It is however perhaps surprising that there is a change at all when it is known that HIF-1 is regulated at the post-translational level. Thus these data are suggesting that RIC is having an effect at the transcriptional level. HIF-2 has a similar function to HIF-1 in endothelial cell responses to hypoxia. There is a switch from HIF-1 to HIF-2 which is part of the cellular adaption to hypoxic stress.^47^ VEGFα is major marker of hypoxic stress, regulated by HIF.^48^

These data together show how RIC is mitigating the hypoxic stress in the tissues. Whether this is part of the protective mechanism of RIC or simply a marker of the reduced injury because of RIC is not clear but it is certainly is plausible that HIF and VEGF are part of a pathway by which RIC is protecting the tissue. Similarly, The cytokeratines examined KRT-7, -8, -18 and -19 all showed the same pattern of an increase with IRI which was mitigated by RIC (Figure 12). This suggest that either they are playing an role in the pathology of IRI that is reduced by RIC leading to less injury or conversely that they are a marker of the severity and requirement for healing of the injury. In this context fibroblasts and smooth muscle markers were increased by IRI with the effect mitigated by RIC (Figure 11). These cells are involved in wound repair and are required to support tissue healing and homestasis to maintain the epithelail barrier and thus maybe protective in NEC pathophysiology.

**Figure 11.**
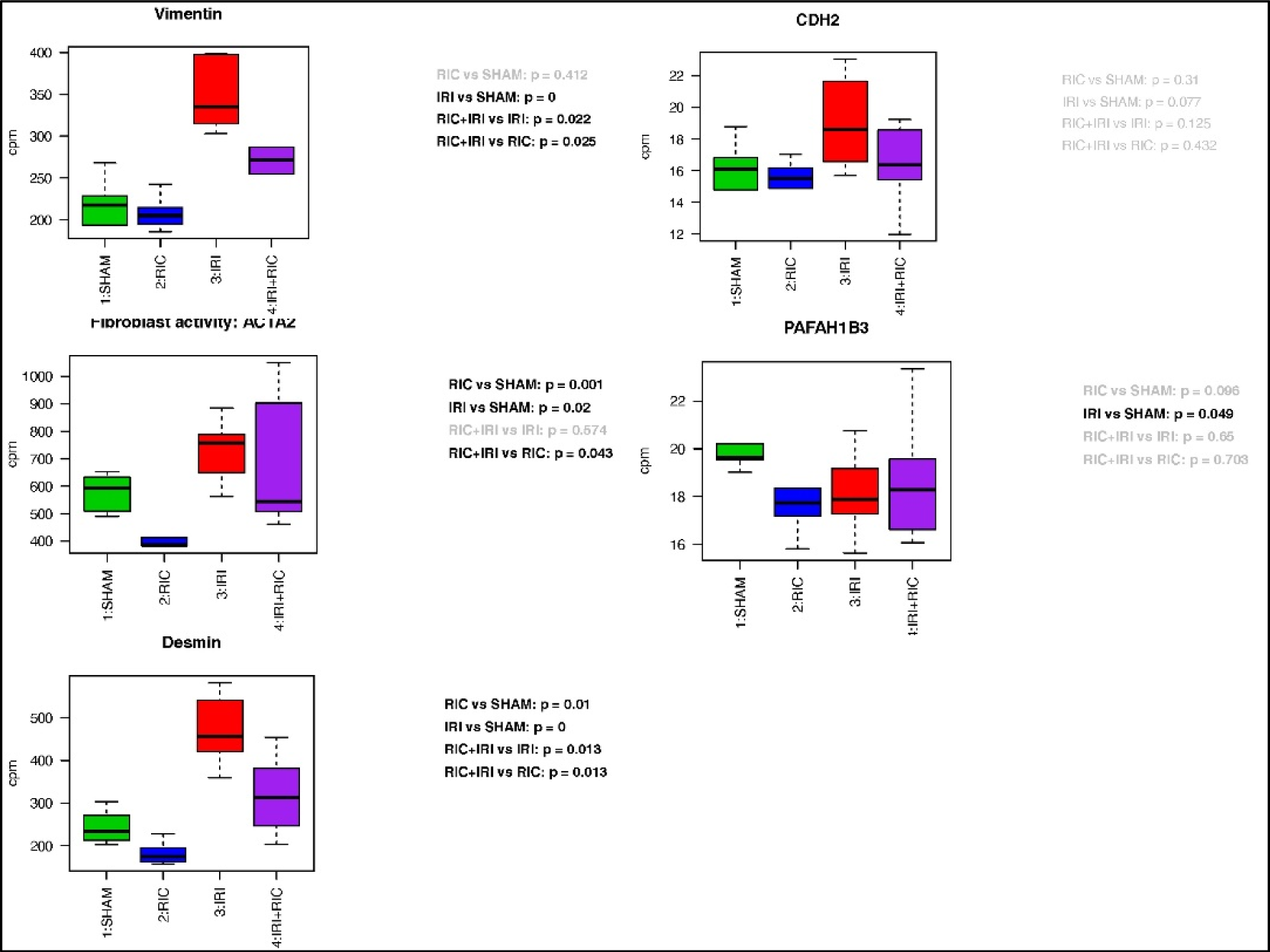
Four way comparison of gene expression of specific markers: Cell type markers for fibroblasts and alpha smooth muscle.

**Figure 12.**
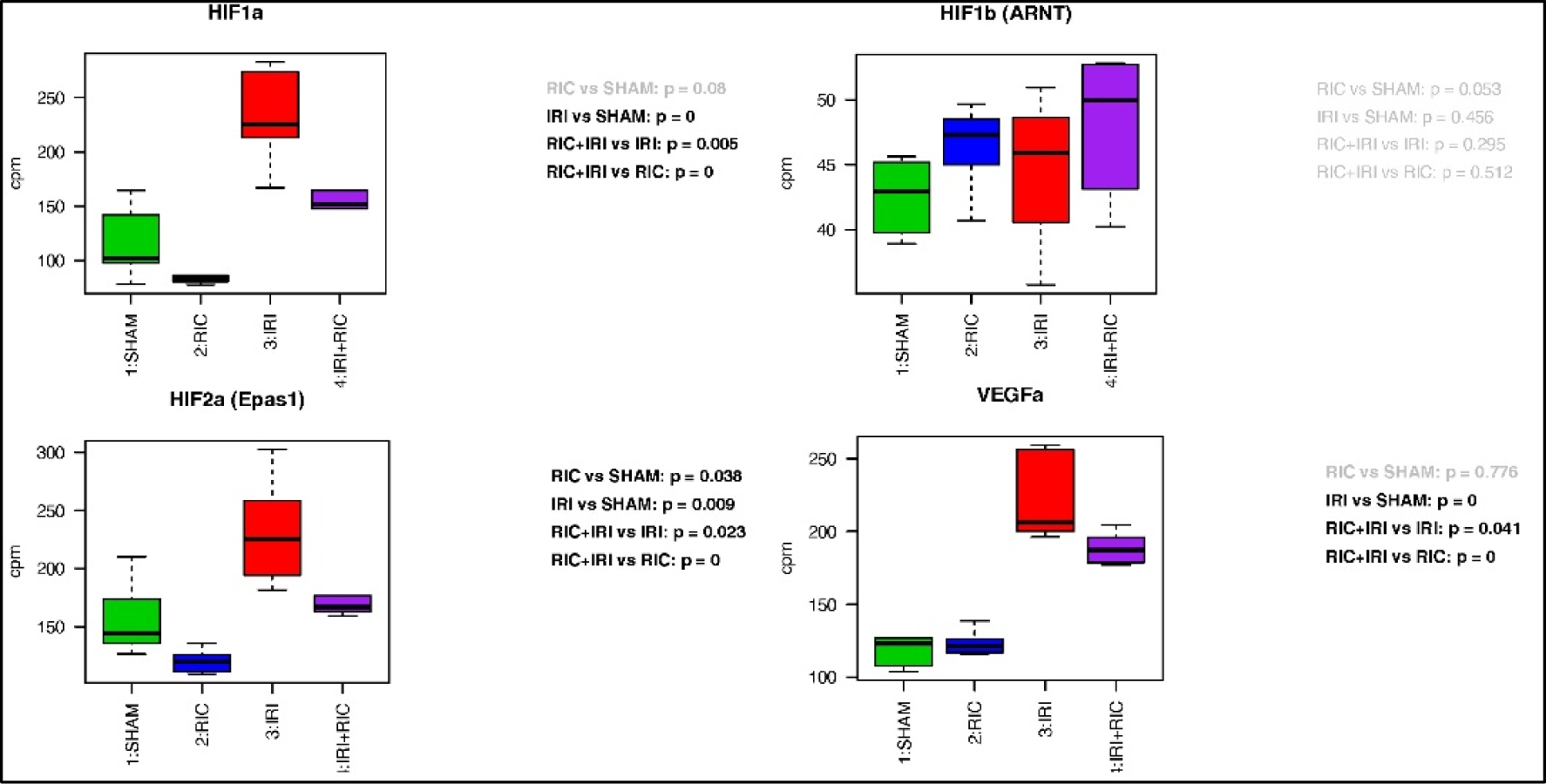
Four way comparison of gene expression of specific markers: Hypoxic pathways.

Toll-like receptor 4 (TLR4) is thought to be a key part of the mechanism. The Toll-like receptors are part of the innate immune system that trigger an inflammatory response to bacterial infection by recognising common epitomes that are highly conserved in bacterial species. TLR4 activation triggers increased expression of MyD88 and NF-ĸβ. These data show that TLR4 expression is decreased by IRI. However, the down-stream effects of increased MyD88 and NF-ĸβ are seen in this model. Moreover, both of these genes show reduced expression in response to RIC. In the case of MyD88, there is a mitigation of the effect of IRI therefore the reduction might not be realised if something other than IRI was triggering the rise. However, NF-ĸβ expression is suppressed independent of IRI.

This supports the concept that the model is a reasonable representation of the human disease and that there is some cross-over in the pathways such that RIC could have a protective effect against NEC both in terms of reducing the necrosis but also in terms of interrupting the pathophysiological pathways earlier. Whilst RIC has of course been investigated primarily as a potential therapy for specifically ischaemic injury, more recent work as looked at the anti-inflammatory potential.^49^ Because these pathways interact and overlap, there is good reason to think that RIC may have a beneficial effect even in the earlier stages of NEC before necrosis develops.

The reperfusion injury salvage kinase pathway (RISK), in the context of cardiac ischaemia-reperfusion is an important part of the protective cellular mechanism. We hypothesise that there is an equivalent pathway in the intestine. Genes in this pathway include MEK1, MEK2, and PI3K. In each case the expression of these genes is decreased by IRI. With MEK1, the expression is unchanged by RIC. With MEK2, RIC decreases the expression but not be as much as IRI does. In the case of PI3K, RIC increases the expression. The RISK pathway is an anti-apoptotic cascade and works by inhibiting the opening of the mitochondrial permeability transition pore (mPTP).^22^ Therefore it is not surprising that in the context of widespread ischaemia all the proteins in the pathways show reduced expression. Equally, because it is a cascade (primarily mediated by phosphorylation), it is also not surprising that there is no clear change in expression seen at the RNA level. A cascade like this works by (in this case) phosphorylation of proteins. Thus a change in the expression level of just one of the proteins involved with have a knock-on effect on all the down-stream proteins.

The increase in expression PI3K, due to RIC, is seen in both the animals exposed to IRI and those that were not, suggesting a potential key role here for this molecule in the protective mechanism of RIC in relation to the RISK pathway. These data show that RIC increases this expression and that if IRI occurs, expression, whilst much lower than baseline is still higher than that seen in IRI alone. IRI results in a decrease in expression but this decrease is mitigated by RIC. Phosphoinositide 3-kinases are a family of signal transducer enzymes that function by phosphorylating the 3 position hydroxyl group of the inositol ring of phosphatidylinositol.^50^ Hausenloy *et al.* (2012) studied the role of P13K in RIC in the porcine heart.^51^ Their data showed that the protective effect of RIC on the heart could be abolished by administration of a blocking agent of PI3K (Wortmannin). The role of PI3K was also confirmed by Western-blotting analysis. The increase in PI3K expression in the intestine demonstrated here would be consistent with an equivalent RISK pathway existing in the intestine. In these data, PI3K increased by RIC (RIC vs SHAM: p = 0.023; RIC+IRI vs IRI: p < 0.001), Table 5, Figure 14.

**Figure 13.**
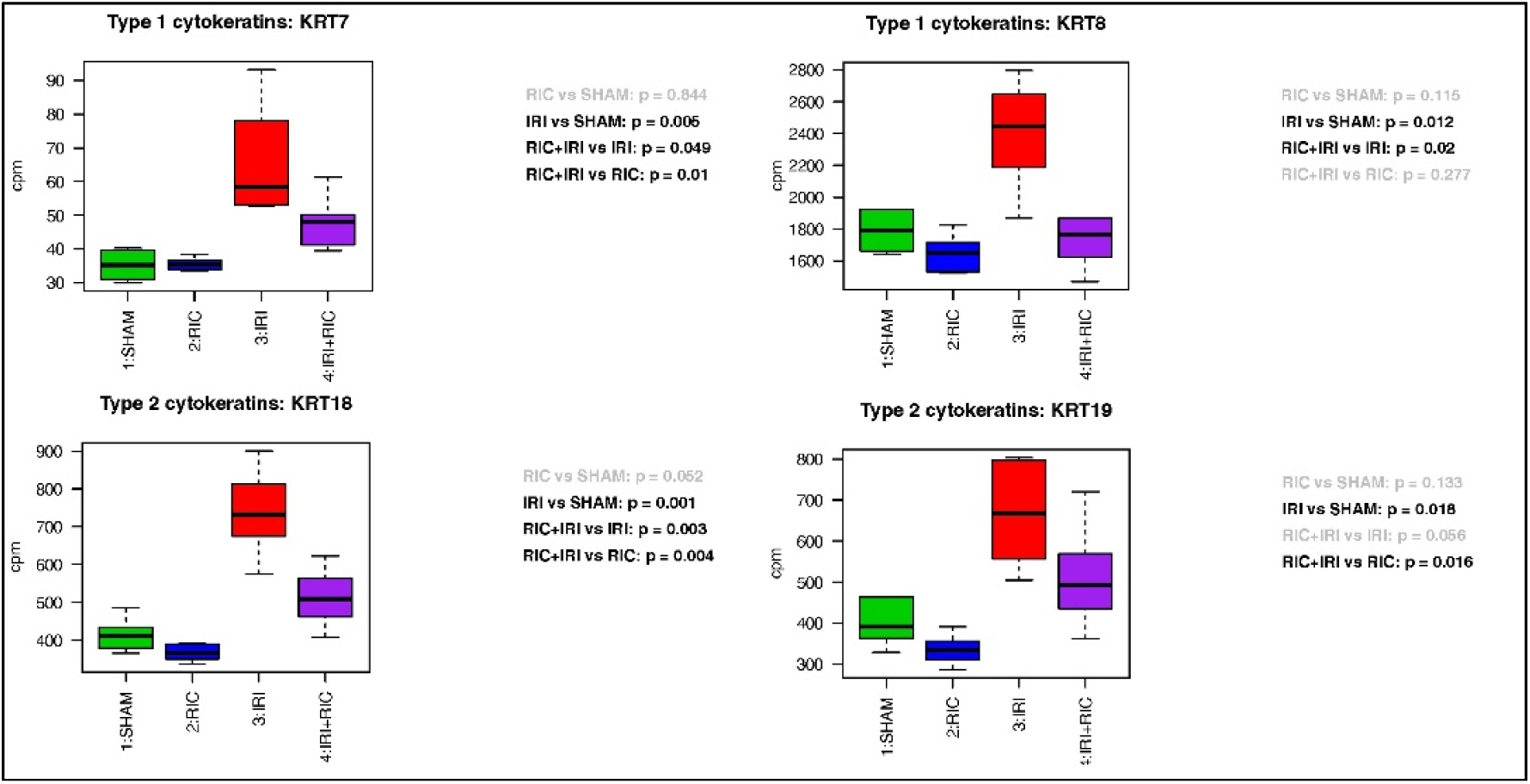
Four way comparison of gene expression of specific markers: markers of cell toxicity.

**Figure 14.**
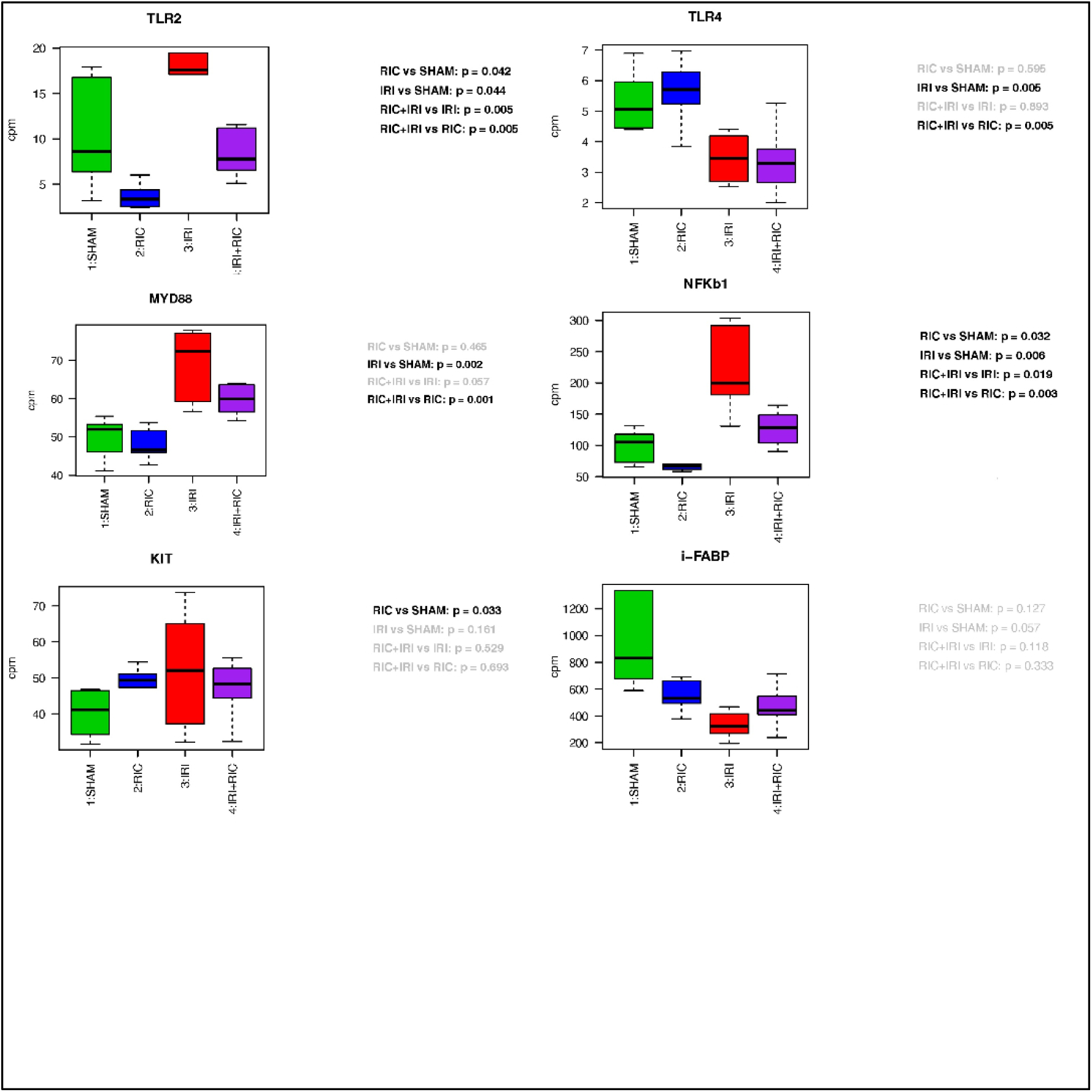
Four way comparison of gene expression of specific markers: Known to be involved in NEC pathophysiology.

P38 and Protein Kinase C were studied by Heinen *et al.* (2011)^52^ with Western Blotting analysis to study the protective pathways of ischaemic conditioning – both direct and remote. Their data showed a role for P38 and PKC in direct ischaemic conditioning but in their data in the heart, the expression of P38 and PKC did not change with *remote* conditioning.^52^ These data are consistent with those results as both P38 (IRI vs SHAM: p = 0.001; RIC+IRI vs RIC; p = 0.012) and PKC (IRI vs SHAM: p < 0.001; RIC+IRI vs RIC, p = 0.01) in the intestine show altered expression with IRI but not with RIC. Table 5, Figure 15.

**Figure 15.**
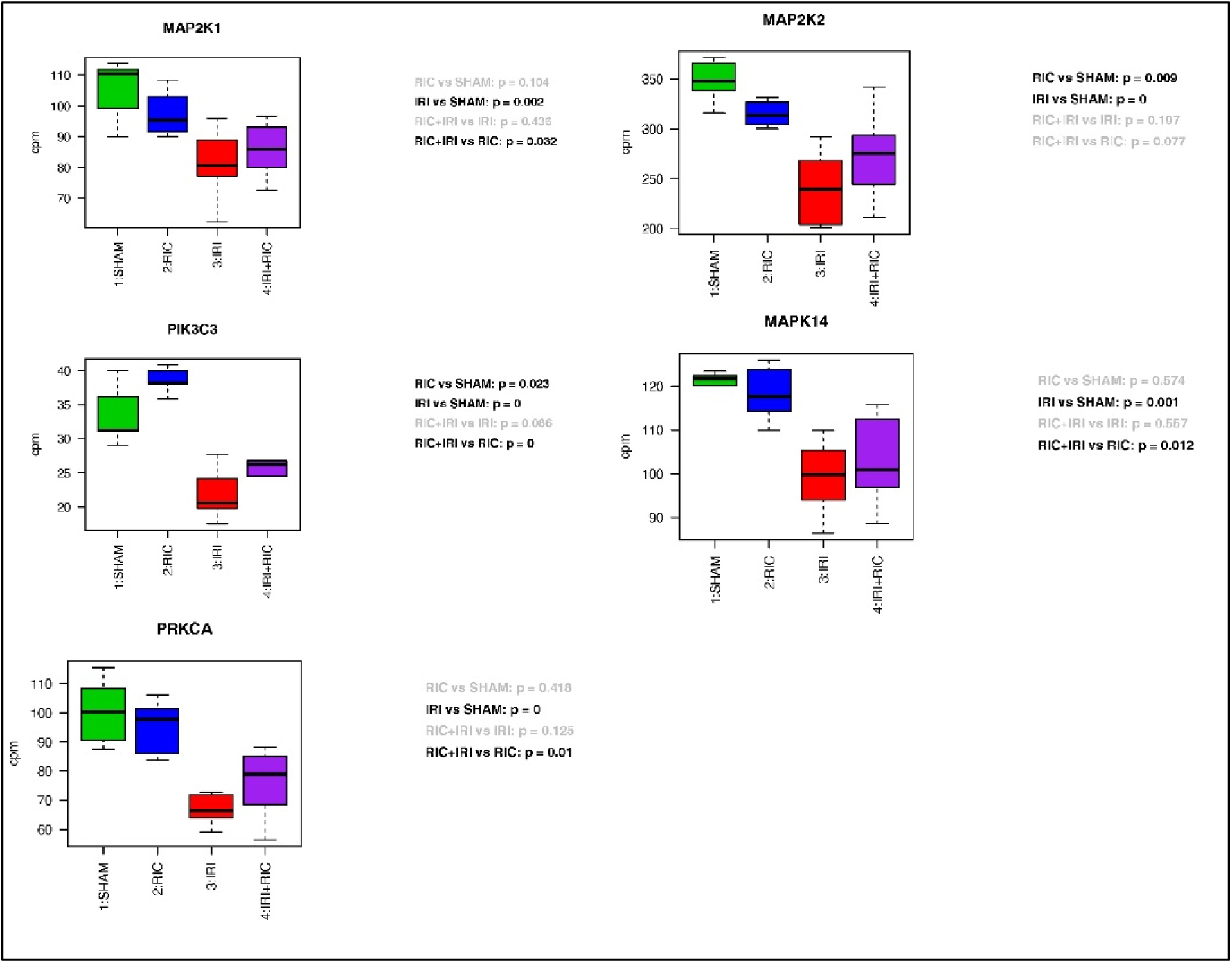
Four way comparison of gene expression of specific markers: Equivalent of RISK pathway.

It is intriguing that these pathways clearly play a role is the protective effect of conditioning when the conditioning is applied directly to the target organ but seem to have little/no role in the same protective effect when the conditioning is delivered remotely. However, these results are entirely consistent with these findings in an entirely different organ system. Further supporting the hypothesis that the mechanisms in the intestine are similar to that that seen in cardiac tissue.

<u>N</u>o previous studies have been published examining the transcriptome of the intestine in response to RIC. Multiple pathways of NEC pathogenesis have been proposed. No animal model completely replicates the human disease and rats (used in this work) are no exception. The importance of bacterial colonisation and invasion in the human disease is certainly not fully replicated in this (or any other) model. However, there are immune pathways that are important in NEC before it gets to the stage of necrosis. This model best mimics the common end-point of NEC with a pattern of bowel necrosis and systemic effects very akin to the human disease. The mechanisms by which RIC is protective may also function at this level – by driving an anti-inflammatory response as well as the resistance to ischaemia; if this is true then RIC would potentially provide a protective effect against NEC earlier in the pathophysiology.

In conclusion, the data shown here suggest several promising avenues for research. The changes in gene expression seen by comparing IRI with the SHAM group show that this model, as well as generating a macroscopic injury that is similar to severe NEC, ^4,8,9^ mimics many known biochemical pathophysiological pathways. There is evidence that an intestinal equivalent of the RISK pathway in cardiac tissue may exist, with data showing significant increases in PI3K with RIC and RIC/IRI and that RIC is triggering an anti-inflammatory process, involving known pathways of the innate immune system, including NFKB, a very important regulator of inflammation.

## Methods

### Animal model^4,8,9^

Animal experiments were carried out with ethical approval from the local Animal Welfare and Ethical Review Body and according to the Animals in Scientific Procedures Act (APSA) 1986 and revisions (Project licence: PA813F125). This study is reported in accordance with the ARRIVE guidelines.^15^ Intestinal injury was induced in suckling rat pups aged 10-13 days, under terminal anaesthesia. Laparotomy was performed and IRI induced by occlusion of the SMA with a microvascular clip. Controls underwent SHAM surgery. RIC was induced by means of a ligature applied to the hind limb to occlude arterial inflow. Each animal undergoing RIC received 3 cycles of 5 minutes of ischaemia with 5 minutes of reperfusion between the ischaemia episodes. The animals were randomly allocated to four groups: SHAM; RIC; IRI; RIC+IRI (Figure 16).

**Figure 16.**
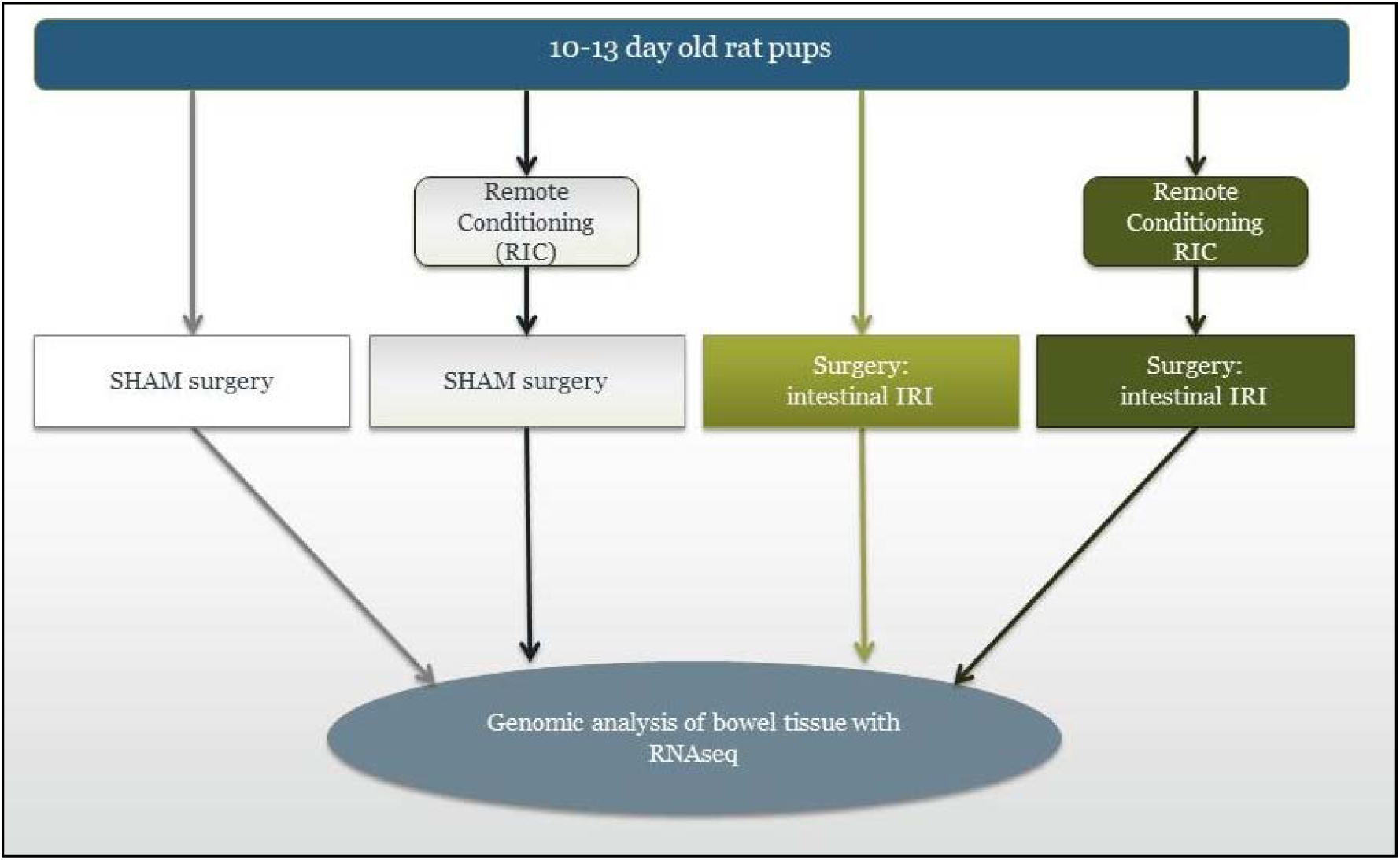
Experimental groups: Related pups were randomly allocated to four groups: Controls that underwent SHAM surgery only (SHAM); animals that underwent hind limb remote ischaemic conditioning and SHAM surgery (RIC); animals that underwent intestinal ischaemia and reperfusion only (IRI); and animals that underwent both RIC and intestinal IRI (RIC+IRI). RNAseq was performed on terminal ileum samples (n=6 for each group).

### RNA isolation and preparation

At the end of the procedure, 2 cm of ileum (3-5 cm from the ileo-caecal valve) was removed for quantitative RNA analysis. This was immediately flushed with Phosphate Buffered Saline (with calcium chloride and magnesium chloride.) (Gibco / Thermo Fisher Scientific) and then placed in RNAlater™ Stabilization Solution (Qiagen) and stored at -20°C, as per the manufacturer’s instructions. The RNA extraction, library preparation and next-generation sequencing were performed by Qiagen Genomic Services (Hilden, Germany). Genome alignment was also performed by Qiagen, who provided the raw-count matrix used for analysis of differential expression. The workflow is shown in Figure 17. The animal experimental protocol is described in full in our previous publication.^4^

**Figure 17.**
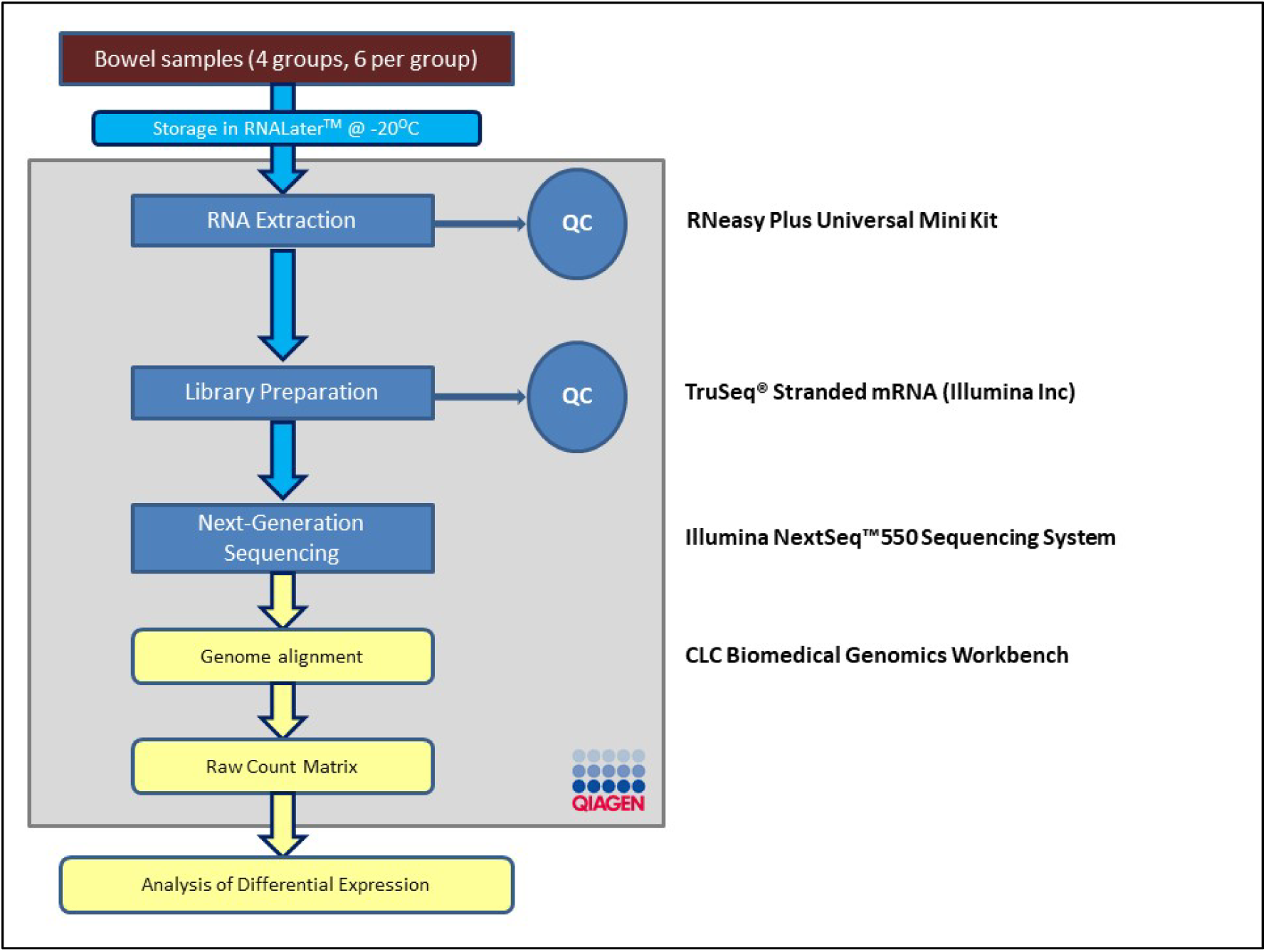
Samples of bowel tissue were sent to Qiagen in RNAlater^TM^. All the steps in the grey box were performed by Qiagen Laboratories. The ‘Wet-lab’ work prior to RNA extraction is described in detail Jones *et al.* (2022).

RNA Extraction and quality control was performed using the RNeasy Plus protocol which uses phenol/guanidine for lysis and silica-membrane purification of RNA. Sample quality control was then performed on each sample. The quantity of RNA was determined spectrophotometrically by measuring the absorbance at 260 nm. The integrity of the total RNA purified was then assayed with the Agilent 4200 TapeStation system (Agilent Technologies Ltd, USA). 2 μl of each sample was used to assess RNA purity generating an RNA Integrity Number (RIN). RIN scores above 8 are considered ‘high quality;’ scores of 5 - 7 indicate some degree of fragmentation.^10,11^

### Library Preparation

The library was prepared using the TruSeq® Stranded mRNA Sample preparation kit (Illumina Inc) which produces a cDNA li brary for whole transcriptome analysis. Briefly, the starting material (500 ng) of total RNA was mRNA enriched using the oligodT bead system. The isolated mRNA was subsequently fragmented using enzymatic fragmentation. Then first strand synthesis and second strand synthesis were performed and the double stranded cDNA was purified (AMPure XP, Beckman Coulter). The cDNA was end repaired, 3’ adenylated and Illumina sequencing adaptors ligated onto the fragments ends, and the library was purified (AMPure XP). The mRNA stranded libraries were pre-amplified with PCR and purified (AMPure XP). The libraries size distribution was validated and quality inspected on a Bioanalyzer 2100 or BioAnalyzer 4200 TapeStation (Agilent Technologies). High quality libraries are pooled in equimolar concentrations based on the Bioanalyzer Smear Analysis tool (Agilent Technologies). The library pool(s) were quantified using qPCR and optimal concentration of the library pool used to generate the clusters on the surface of a flowcell before sequencing.

### Next-generation Sequencing

The sequencing of the RNA was preformed using an Illumina NextSeq 550 High Throughput Next Generation Sequencer. (Qiagen, Germany / Illumina, USA). The transcriptome was sequenced using the following parameters: 30M reads and a read depth of 75 base pairs.

### Genome Alignment

The genome alignment step of the analysis was performed using CLC read mapper within Qiagen’s Biomedical Genomics Workbench software (Qiagen, Germany / CLC bio, Denmark).^12^ Benchmarking studies have shown this gives equivalent or better performance compared to widely used aligning tools. ^13^

### Analysis

Analysis of the mapped counts was conducted using *R* v 3.6.1 (R Foundation for Statistical Computing, Vienna, Austria).^16^ The *R* Script used is included in the additional material. To test for potential confounders, known phenotypic data for each animal was entered. The age in days, gender, weight, from which litter the animal came and the date of the experiment was entered for each animal and linked to the corrosponding expression values. Before formal analysis of the data, the raw data was explored with principal component analysis (PCA) to perform quality control, with searchs for batch effects and removal of significant outliers from the model if necessary.

### Data Exploration and Quality Control

Lowly expressed mRNA genes were excluded in order to reduce the need for multiple testing correction and thus reducing the risk of a Type II error and inconsistent reads across samples. Using the whole dataset; Hierarchical clustering was performed using the Euclidean distance method; PCA was performed and plotted; and median vs interquartile range for each raw count was plotted. Finally the PCA was plotted with the samples labelled for the known phenotypical factors. Any sample that was shown to be an outlier in more than one of these plots would be considered as a true outlier and excluded from further analysis.

### Differential Expression Testing

The differential gene expression was assessed in four 2-way comparisons, using edgeR (version 3.6.1).^11,12^

### Differential Expression – Targeted Gene Analysis

As well as performing this undirected differential expression testing, we interrogated the data for specific genes of interest. These broke down to various sub-groups; Markers of Cell types (Table 1), markers of biological processes (Table 2), proteins known to be important in the pathogenesis of NEC (Table 3) and proteins that are found in known biological pathways of RIC in other tissues (Table 4). In order to compare these different genes, simple box-plots and t-tests were performed using R.

