## Supplementary material for "Transcriptomic Analysis of the Effect of Remote Ischaemic Conditioning in an Animal Model of Necrotising Enterocolitis": Gene Expression Comparison (t-Tst).R

```
#library(FSA)

setwd("~Files/_PhD/Data/RNA/Processing/Biased analysis")

data<-read.table("Gene Counts_IJ2.csv", sep=',', header=TRUE, row.names=1, check.names=FALSE)

#Testing in same order

group<-data[46,]

boxplot_data<-data[1:45,]

table(names(group) == colnames(boxplot_data))

#boxplots

pdf("./Boxplots_for_individual_genes (t-Tests).pdf")
tTsts_columnnames <-c ("Gene name",  "SHAM vs RIC, p =", "SHAM vs IRI p =","IRI vs RIC+IRI p =", "IRI vs RIC+IRI p =")
tTsts_results_df <- tTsts_columnnames
par(mfrow=c(2,2))


for (i in row.names(boxplot_data)) {
    
  kdata <- as.numeric(boxplot_data[i,])
  SHAM <- kdata [1:6]
  RIC <- kdata [7:12]
  IRI <- kdata [13:18]
  RICIRI <-kdata [19:24]
  RvS <- t.test(RIC, SHAM, alternative = "two.sided", var.equal = FALSE)
  IvS <- t.test(IRI, SHAM, alternative = "two.sided", var.equal = FALSE)
  RIvI <- t.test(RICIRI, IRI, alternative = "two.sided", var.equal = FALSE)
  RIvR <- t.test(RICIRI, RIC, alternative = "two.sided", var.equal = FALSE)
  boxplot(as.numeric(boxplot_data[i,]) ~as.factor(data[46,]), col=c("green3","blue", "red","purple"), outline=FALSE, main=paste(i), las=2,xlab="", ylab="cpm")
  
  tTsts_matrix <- (c(i, RvS$p.value, IvS$p.value, RIvI$p.value, RIvR$p.value))
  
  tTsts_results_df <- rbind(tTsts_results_df, tTsts_matrix)
  
  Comp1 <- paste0("RIC vs SHAM: p = ", round(RvS$p.value,4))
  Comp2 <- paste0("IRI vs SHAM: p = ", round(IvS$p.value,4))
  Comp3 <- paste0("RIC+IRI vs IRI: p = ", round(RIvI$p.value,4))
  Comp4 <- paste0("RIC+IRI vs RIC: p = ", round(RIvR$p.value,4))
  
  plot(NA, xlim=c(0,20), ylim=c(0,4), bty='n',
       xaxt='n', yaxt='n', xlab='', ylab='')
  #text(1,4,kwtext, pos=4, cex=1)
  text(1,3.5,Comp1, pos=4, cex=1)
  text(1,3,Comp2, pos=4, cex=1)
  text(1,2.5,Comp3, pos=4, cex=1)
  text(1,2,Comp4, pos=4, cex=1)
} 
dev.off()


write.csv (tTsts_results_df, "./t_Test_results_for_individual_genes.csv", row.names = FALSE)
```
