## Supplementary material for "Transcriptomic Analysis of the Effect of Remote Ischaemic Conditioning in an Animal Model of Necrotising Enterocolitis": R_code_gene_expression.R

```
#13th November 2019
#Ashley Heinson <>


############################# STARTING SCRIPT FROM HERE TIDY UP.
#Install.packages....

#edgeR nor EnhancedVolcano won't install with install.packages#
#if (!requireNamespace("BiocManager", quietly = TRUE))
 # install.packages("BiocManager")

#BiocManager::install("edgeR")

### Setting WD and reading in files.

#Loading packages at the start.
library(EnhancedVolcano) #not available for 3.6.1
library(gplots)
library(amap) #- amap won't run on R3.5.1 - without amap can't do pearson
#library(dendextend) #probably not needed at this point
library(RColorBrewer)
library(edgeR)
library (biomaRt)

#citations

citation ("EnhancedVolcano")
citation ("gplots")
citation ("amap")
citation ("dendextend")
citation ("RColorBrewer")
citation ("edgeR")
citation (package="biomaRt")


pheno<-read.table("Sample phenotype.csv", header=TRUE, row.names=1, sep=',')
counts<-read.table("Raw_Counts_All_Samples.csv", header=TRUE, sep=',', row.names=1,check.names = FALSE)
#Reads in raw counts and the phenotype information

# Exploring the Raw data (QC) 

# Merging Pheno and Count data together to keep only samples that we have both for and make sure that the samples are in the same order within the files. 
t_counts<-t(counts)
combined<-merge(pheno, t_counts,by="row.names", all=FALSE)
dim(t_counts)
dim(pheno)
dim(combined)

row.names(combined)<-combined$Row.names
combined<-combined[,-1]

save(combined,file="Counts_pheno_combined.RData")


#########################################
#
#
# Starting from here read in combined (load above saved file)
# Data merged in "Counts_pheno_combined"
#
#
###########################################

load("Counts_pheno_combined.RData")

values<-combined[,12:length(colnames(combined))] #takes gene counts in 'values'
info<-combined[,1:8] #takes phenotypic info into 'info'

# Boxplot
pdf("Raw_Boxplot_mRNA_RNA_Counts.pdf")
par(cex=0.3, mar=c(25,10,10,15))
boxplot(t(values), outline=FALSE, las=2, col="gray85", main="Raw Sample Counts mRNA Hits", cex.main=5, label=row.names(values))
boxplot(t(values), outline=TRUE, las=2, col="gray85", main="Raw Sample Counts mRNA Hits", cex.main=5, label=row.names(values))
#raw counts whiskers are 1.5xIQR
dev.off()


##### THERE ARE ALOT OF LOWLY EXPRESSED SMALL RNAs
# Trying filtering out lowly expressed genes just to see if this changes the plots.
# Removing small RNAs that are not expressed in at least half of all samples (45)
values<-t(values)
Filter <-6 #limit of a gene being expressed in how many samples... random number to remove of v.v.lowly expressed genes
#this is artbitary can be stricter or less strict.
# if filter is 6: 18040  - if 10: 17154 - if 2: 19251
keep_tags<-rowSums(values>1)>=Filter
values_filtered<-values[keep_tags,]
#save(values_filtered, file="Raw_Filtered_mRNA_All_Samples.RData")
dim(values)
#dim(values)
#[1] 30967    24
print (Filter)
dim(values_filtered)
#dim(values_filtered)
#[1] 18040   24
#### PLOTTING FILTERED SAMPLES
# Boxplot
pdf("Raw_Filtered_Boxplot_mRNA_Only_RNA_Counts.pdf")
par(cex=0.3,mar=c(25,10,10,15))
boxplot(values_filtered, outline=FALSE, las=2, col="gray85", main="Raw Filtered Sample Counts mRNA Hits", cex.main=5)
#Same as last charts with very lowly expressed removed.
dev.off()


#Hierachical Clustering
values_filtered<-t(values_filtered)


distance_euc<-dist(values_filtered,method="euclidean")
#distance_pear<-dist(values_filtered,method="pearson")

hc_euc_av<-hclust(distance_euc, method="average")
hc_euc_comp<-hclust(distance_euc, method="complete")
#hc_pear_av<-hclust(distance_pear, method="average")
#hc_pear_comp<-hclust(distance_pear, method="complete")
hc_euc_ward.D<-hclust(distance_euc, method="ward.D")
#hc_pear_ward<-hclust(distance_pear, method="ward")


pdf("Clustering_Raw_mRNA_All_Data_Filtered.pdf", w=18)
par(cex=0.5)
plot(hc_euc_av, main="genes_filtered Euclidean Average", hang=-1)
plot(hc_euc_comp, main="genes_filtered Euclidean Complete", hang=-1)
#plot(hc_euc_ward, main="genes_filtered Euclidean Ward", hang=-1)
#plot(hc_pear_av, main="genes_filtered Pearson Average", hang=-1)
#plot(hc_pear_comp, main="genes_filtered Pearson Complete", hang=-1)
#plot(hc_pear_ward, main="genes_filtered Pearson Ward", hang=-1)
dev.off()

#PCA Analysis
#library(rgl)
pcs<-prcomp(values_filtered)
PC1<-pcs$x[,1]
PC2<-pcs$x[,2]
PC3<-pcs$x[,3]
PCA_details<-cbind(PC1,PC2,PC3)
pca<-merge(PCA_details,info,by="row.names",all=FALSE)

row.names(pca)<-pca$Row.names
pca<-pca[,-1]

PC1<-pca$PC1
PC2<-pca$PC2
PC3<-pca$PC3

plot(PC1,PC2,main="PCA Raw Counts RNA All Data")

pdf("PCA_Raw_Counts_mRNA_All_Data_Filtered.pdf")
#par(mfrow=c(3,2))
plot(PC1,PC2,main="PCA Raw Counts RNA All Data")
plot(PC1,PC2,main="PCA Raw Counts RNA All Data")
text(PC1,PC2,labels=row.names(pca),cex=0.3)
dev.off()

#IQR/Median Plotting
# Generating IQR and Median plots
# margin 1 = rows, 2 = columns
IQR_samples<- apply(values_filtered, 1, IQR)
Median_samples<- apply(values_filtered, 1, median)

SD_IQR<-sd(IQR_samples)
SD_Median<-sd(Median_samples)
Mean_IQR<-mean(IQR_samples)
Mean_Median<-mean(Median_samples)
SD_1_IQR<-c((Mean_IQR - SD_IQR), (Mean_IQR + SD_IQR))
SD_1_Median<-c((Mean_Median - SD_Median), (Mean_Median + SD_Median))

SD_2_IQR<-c((Mean_IQR - (2 * SD_IQR)), (Mean_IQR + (2 *SD_IQR)))
SD_2_Median<-c((Mean_Median - (2 * SD_Median)), (Mean_Median + (2 * SD_Median)))

pdf("IQR_RAW_mRNA_All_Samples_Filtered.pdf")
plot(Median_samples, IQR_samples, main="IQR Median Plot of the Raw Counts mRNA All Samples", xlim=c(300,420),ylim=c(1000,1300))
points(SD_1_Median, SD_1_IQR, pch=3, col="firebrick1", cex=1)
points(SD_2_Median, SD_2_IQR, pch=3, col="firebrick3", cex=1)

plot(Median_samples, IQR_samples, main="IQR Median Plot of the Raw Counts mRNA All Samples",xlim=c(300,420),ylim=c(1000,1300))
points(SD_1_Median, SD_1_IQR, pch=3, col="firebrick1", cex=1)
points(SD_2_Median, SD_2_IQR, pch=3, col="firebrick3", cex=1)

text(Median_samples,IQR_samples,labels=row.names(values_filtered),cex=0.3)


dev.off()

#### ONLY NEEDED IF REMOVING OUTLIERS

##### Outlier Removal:
#outliers<-c("SampleID_22.H.AN")
# Removing the outliers from filtered Matrix
#values_filtered<-t(values_filtered)
#values_filt_out_rem <- values_filtered[!row.names(values_filtered) %in% outliers,]
#info_out_rem<-info[!row.names(info) %in% outliers,]
#save(values_filt_out_rem, file="./Data/Raw_Filtered_Out_Removed_mRNA_Only_RNA_All_Samples_Macrophages_Exosomes_Applied.RData")
#write.table(info_out_rem, file="./Data/Phenotype_Data_Outliers_Removed_mRNA_Only_Final_Macrophages_Exosomes_Applied.csv", sep=',', col.names=NA)

##### PCA Plot to explore the variation of the datasets:
### ONLY NEED TO REDO PLOTS ONLY IF OUTLIERS ARE REMOVED
#pcs<-prcomp(values_filt_out_rem)
#PC1<-pcs$x[,1]
#PC2<-pcs$x[,2]
#PC3<-pcs$x[,3]
#PCA_details<-cbind(PC1,PC2,PC3)
#write.table(PCA_details, "./Data/PCA_Raw_Filtered_NEW_Outlier_Removed_mRNA_Only_RNA.csv", sep=",")

# Add Demo Data for colouring. 
#pca <-  read.table("./Data/PCA_Raw_Filtered_NEW_Outlier_Removed_mRNA_Only_RNA.csv",header=TRUE, sep=",",row.names=1)
#PC1<-pca$PC1
#PC2<-pca$PC2
#PC3<-pca$PC3

### Plotting As A Continous Colour Palette Legend is Min and Max
# Optionally set colours using RColorBrewer

cols = brewer.pal(4, "Blues")
#To view the colorbrewer palletes in R:
#display.brewer.all(5)

# Define colour pallete
#pal = colorRampPalette(c("blue", "red"))
# Use the following line with RColorBrewer
pal = colorRampPalette(cols)

pdf("Colour_coded_for_known_variables.pdf")
par(mfrow=c(3,2))

for (i in 6:length(colnames(pca))){
  print(i)
  print(colnames(pca[i]))
  pca_current<-pca
  # Rank variable for colour assignment
  pca_current$order = findInterval(as.numeric(pca_current[,i]), sort(as.numeric(pca_current[,i])))
  
  # Make plot
  plot(PC1,PC2, pch=19, col=pal(nrow(pca_current))[pca_current$order], main=paste("PCA Coloured By ",colnames(pca[i]),sep=""))
  # Add a simple legend
  legend("topleft", col=pal(2), pch=19,
         legend=c(round(range(as.numeric(pca_current[,i])), 1)))
  rm(pca_current)
}

dev.off()


plot3d(PC1,PC2,PC3, xlab="PC1", ylab="PC2", zlab="PC3", box=FALSE, col=as.integer(pca$Condition), type='s', size=1)
plot(PC1,PC2,main="PCA Raw Counts Small RNA All Data", col=as.factor(pca$Group))
text(PC1,PC2,label=pca$Donor)

pdf("PCA_2D_Raw_Filtered_Counts_mRNA_All_Data.pdf")
par(mfrow=c(2,2))

for (i in 6:length(colnames(pca))){
  print(i)
  print(colnames(pca[i]))
  plot(PC1,PC2, main=paste("PCA Coloured By ",colnames(pca[i]),sep=""), col=as.factor(pca[,i]))
}
dev.off()  

pdf("PCA_2D_Raw_Filtered_Counts_mRNA_All_Data_TEXT.pdf")
for (i in 6:length(colnames(pca))){
  print(i)
  print(colnames(pca[i]))
  plot(PC1,PC2, main=paste("PCA Coloured By ",colnames(pca[i]),sep=""), col=as.factor(pca[,i]))
  text(PC1,PC2,labels=pca[,i],cex=0.8)
  
}
dev.off()  


#### Variances explained by the Principal Components:
test<-data.frame(summary(pcs)$importance)
prop<-test[3,]

pdf("PCA_Variance_Explained_Filtered_Out_Rem_mRNA.pdf")
barplot(as.numeric(prop),names.arg = colnames(prop), las=2, cex.names=0.7, xlab="Principal Components", ylab="Percentage of Variance Explained", main="PCA Percentage of Variance Explained")
dev.off()
write.table(prop, file="PCA_Variance_Explained_Filtered_Out_Rem_mRNA.csv", sep=',')


#########################################
#
#
# Starting from here read in combined (load above saved file)
# Data merged in "Raw_Filtered-mRNA_All_samples.RData"
#
#
###########################################

load("Raw_Filtered_mRNA_All_Samples.RData")
pheno<-read.table("Sample phenotype.csv",row.names=1, sep=',', header=TRUE)

#Save each analysis in its own subfolder - create the subfolder ID to access later
#These subfolders need to be created in the workingdir
#'subdir' is a vector of the four subdirectory names relative to the working directory
subdir <-c("./results_IRI_vs_SHAM/",
           "./results_RIC_vs_SHAM/",
           "./results_RIC+IRI_vs_IRI/",
           "./results_RIC+IRI_vs_SHAM/",
           "./results_RIC+IRI_vs_RIC/"
           )

values_filtered<-t(values_filtered)
combined<-merge(pheno, values_filtered, by="row.names",all=FALSE)
row.names(combined)<-combined$Row.names
combined<-combined[,-1]
values_filtered<-combined[,12:length(colnames(combined))]
pheno<-combined[,1:8]


row.names(values_filtered) == row.names(pheno)


#Set up the experimental Design
#Covariate Model

Condition<-as.factor(pheno$Group)
#OTHERFACTOR<-as.factor(pheno$factor)


design<-model.matrix(~0+Condition) #OTHERFACTOR: condition+OTHERFACTOR

#Read values (genes) into edgeR.
eds<-t(values_filtered)
eds_filtered<-DGEList(counts=eds, genes=rownames(eds))
#Creates rownames required by EdgeR

### NORMALIZE THE DATA
#Account for different library sizes etc.
eds_filtered<-calcNormFactors(eds_filtered,method="TMM")

#Calculating Differentially Expressed Genes:
eds_filtered<- estimateDisp(eds_filtered, design)

# Saving a normalized / filtered cpm matrix
counts.per.m <- cpm(eds_filtered, normalized.lib.sizes=TRUE)

#export Normalized counts table to a file 
write.table(counts.per.m, file="filtered_matrix_counts_per_million.csv", sep = ",", col.names=NA)
#create the matricies of counts per million for the 2-way comparisons:
IRI<-counts.per.m[,1:6]
RIC<-counts.per.m[,7:12]
RICIRI<-counts.per.m[,13:18]
SHAM<-counts.per.m[,19:24]
IRIvsSHAM<-cbind(IRI,SHAM)
RICIRIvsIRI<-cbind(RICIRI,IRI)
RICvsSHAM<-cbind(RIC,SHAM)
RICIRIvsSHAM<-cbind(RICIRI,SHAM)
RICIRIvsRIC<-cbind(RICIRI,RIC)
rm(IRI)
rm(RIC)
rm(SHAM)
rm(RICIRI)


comp<-c("IRI_vs_SHAM",
        "RIC_vs_SHAM",
        "RIC_and_IRI_vs_IRI",
        "RIC_and_IRI_vs_SHAM",
        "RIC_and_IRI_vs_RIC"
) #Filenames for 5 groups in each of the subfolders

#for test runs of the loop:
#     IRI vs SHAM = 1
#     RIC vs SHAM = 2
#     RIC and IRI vs IRI = 3
#     RIC and IRI vs SHAM = 4
#     RIC and IRI vs RIC = 5

#write these five matricies to csv files
write.table(IRIvsSHAM, file=(paste0(subdir[1],"filtered_matrix_counts_per_million_",comp[1],".csv")), sep = ",", col.names=NA)
write.table(RICvsSHAM, file=(paste0(subdir[2],"filtered_matrix_counts_per_million_",comp[2],".csv")), sep = ",", col.names=NA)
write.table(RICIRIvsIRI, file=(paste0(subdir[3],"filtered_matrix_counts_per_million_",comp[3],".csv")), sep = ",", col.names=NA)
write.table(RICIRIvsSHAM, file=(paste0(subdir[4],"filtered_matrix_counts_per_million_",comp[4],".csv")), sep = ",", col.names=NA)
write.table(RICIRIvsRIC, file=(paste0(subdir[5],"filtered_matrix_counts_per_million_",comp[5],".csv")), sep = ",", col.names=NA)


fit <- glmQLFit(eds_filtered, design)
#Changes column names
colnames(design)<-c("ConditionIRI","ConditionRIC","ConditionRICIRI","ConditionSHAM")

my.comparisons<-makeContrasts(
  IRI_vs_SHAM = ConditionIRI - ConditionSHAM,
  RIC_vs_SHAM = ConditionRIC - ConditionSHAM,
  RIC_and_IRI_vs_IRI = ConditionRICIRI - ConditionIRI,
  RIC_and_IRI_vs_SHAM = ConditionRICIRI - ConditionSHAM,
  RIC_and_IRI_vs_RIC = ConditionRICIRI - ConditionRIC,
  levels=design)

filteredresults<-paste0(subdir,"filtered_matrix_counts_per_million_",comp,".csv") #creates the vector to open each the data for each comparison
readtbls<-paste0(subdir,comp,"_DE.csv") # this selects only differentially expressed genes for heatmap
readtbls_topx<-paste0(subdir,comp,"_t50d_t50U.csv") #this selects only top x of differentially expressed genes for heatmap


#filteredresults<-c("filtered_matrix_counts_per_million_IRI_vs_SHAM.csv","filtered_matrix_counts_per_million_RIC_vs_SHAM.csv","filtered_matrix_counts_per_million_IRI_vs_IRIRIC.csv","filtered_matrix_counts_per_million_IRI_vs_SHAM.csv")
#readtbls<-c("IRI_vs_SHAM_DE.csv","RIC_vs_SHAM_DE.csv","RIC_and_IRI_vs_IRI_DE.csv","RIC_and_IRI_vs_SHAM_DE.csv")
#NEED TO CHECK THESE ARE ALL CORRECT!


for (i in 1:5) {
    qlf <- glmQLFTest(fit, contrast=my.comparisons[,comp[i]])
    #Writing out all genes so that I can pull out log FC and convert to gene symbol for NPA
    all_genes<-as.data.frame(topTags(qlf, n=nrow(eds_filtered)))
    #Sort genes based on their significance and then pull the counts per million expressi n level data from eds_filtered for this ordered list.
    all_results<-(all_genes)
    #Extract only the genes differentially expressed with an FDR corrected p-value <0.05
    DE_results<-all_results[all_results[,"FDR"]<=0.05,]
    #Nom_sig<-all_results[all_results[,"PValue"]<=0.01,]
  
    #Add gene names to DE_results file
    ensembl = useEnsembl(biomart="ensembl", dataset="rnorvegicus_gene_ensembl", mirror = "uswest") # load rat genome
    mart <- useMart(biomart = "ensembl", dataset = "rnorvegicus_gene_ensembl") #mart
    gene.name.search.matrix <-as.matrix(row.names(DE_results))
    SearchList = gene.name.search.matrix
    
    gene.name<-getBM(attributes = c("ensembl_gene_id","external_gene_name"),
                   filters = "ensembl_gene_id",
                   values = SearchList,
                   mart = ensembl) 
   
    DE_results <- cbind(DE_results,gene.name)
   
    
    
    
    
    
    ### Save the differential testing - ALL genes and only the significant ones (miRNAs in this case).
    #write.table(Nom_sig, file="./Results/DEG_res_NOM_SIG_Block_Design_Tumour_Healthy.csv", sep=",", col.names=NA)
    
    fn<-paste0 (subdir[i],comp[i],"_DE.csv")
    write.table(DE_results, file=fn, sep=",", col.names=NA)
    fn<-paste0 (subdir[i],comp[i],"_all_results.csv")
    write.table(all_results, file=fn, sep=",", col.names=NA)

  ## Volcano Plots

  #colnames(all_results)
  #Standard volcano
  data<-all_results  
  data$Colour<-"black"
  data$Colour[abs(data$logFC) > 1 & data$FDR < 0.05]="firebrick2"
  data$Colour[abs(data$logFC) < 1 & data$FDR < 0.05]="darkorange1"
  fnp<-paste0(subdir[i],comp[i],"_Volcano_plots.pdf")
  pdf(fnp)
  mtitle<-paste(comp[i],"volcano plot")
  plot(data$logFC, -log2(data$FDR), ylim=c(0,20), xlab="Log Fold Change", 
      ylab="Negative Log2 False Discovery Rate Corrected P Value", main=mtitle,
      col = data$Colour, pch=20)
  legend("topleft", c("< 0.05 Q Val", "< 0.05 P Val and > 1logFC"), col=c("darkorange1", "firebrick2"), pch=c(20,20))
  dev.off()
  #Enhanced volcano
  fnp<-paste0(subdir[i],comp[i],"_Enhanced_Volcano_plots.pdf")
  pdf(fnp)
  mtitle<-paste(comp[i],"enhanced volcano plot")
  EnhancedVolcano(data, 
                  lab=NA,
                  x='logFC',
                  y='FDR',
                  ylim = c(0,10),
                  title = mtitle,
                  pCutoff= 0.05,
                  FCcutoff=2,
                  pointSize = 1.5,
                  labSize = 1
  )
  dev.off()
  #Reorder differential expression by logFC
  
  fn<-paste0(subdir[i],comp[i], "_DE.csv")
  
  dat<-read.csv(fn, header=T, sep=",") # opens differential expression file
  #dat<-dat[,2:7] #removes duplicated column
  dat2 <-dat[order(dat$logFC),] #reorders according to logFc  
  neg <- dat2$logFC <0 #downregulated genes only
  pos <- dat2$logFC >0 #upregulated genes only
  datneg <- dat2[neg,] #creates dataframe of downregulated genes (biggest change at the top)
  datpos <- dat2[pos,] # creates dataframe of upregulated genes
  datpos <-datpos[order(datpos$logFC,decreasing=TRUE),] #reorders upregulated so biggest change at the top)
  fnU<-paste0(subdir[i],comp[i], "_DE_UpReg.csv")
  fnD<-paste0(subdir[i],comp[i], "_DE_DwnReg.csv")
  fnC<-paste0(subdir[i],comp[i],"_t50D_t50U.csv") #change this to make more accurate
  
  write.csv(datpos,fnU) #Up regulated genes
  write.csv(datneg,fnD) #Down regulated genes
  
  #Selection of most DE gene subset.
  #Set the value of x for the number of pos AND neg genes (i.e. total = 2x)
  x <-25 #change this number for number of selected genes i.e. 50 most neg and 50 most pos changed
  #because there might not be enough down or up regulated genes:
  neg_count<-nrow(datneg)
  if (neg_count > x) {neg_count<-x}
  pos_count<-nrow(datpos)
  if (pos_count > x) {pos_count<-x}
  
  datcombined <-rbind(datpos[1:pos_count,],datneg[1:neg_count,]) #top x up and down regulated 
  write.csv(datcombined,fnC) #Top x of both up and down regulated
  
  
  
  #Heatmaps

  c<-rep(1:500,each=1)
  cpms<-read.table(filteredresults[i], header=TRUE, row.names=1, sep=',')
  DeGene<-read.table(readtbls[i], header = TRUE, row.names = 1, sep=",")
  TopGene<-read.table(readtbls_topx[i],header=TRUE, row.names=2, sep=",")
  matrix<-as.matrix(cpms[row.names(cpms) %in% row.names(DeGene),]) #Takes the cpms for differentially expressed genes
  gene.name.search.matrix <-as.matrix(row.names(matrix))
  SearchList = gene.name.search.matrix
  gene.name<-getBM(attributes = c("ensembl_gene_id","external_gene_name"),
                   filters = "ensembl_gene_id",
                   values = SearchList,
                   mart = ensembl) 
  row.names(matrix) <- gene.name[,2]
  
  matrix_topx<-as.matrix(cpms[row.names(cpms) %in% row.names(TopGene),]) #Takes the cpms for most differntially expressed genes
  gene.name.search.matrix <-as.matrix(row.names(matrix_topx))
  SearchList = gene.name.search.matrix
  gene.name<-getBM(attributes = c("ensembl_gene_id","external_gene_name"),
                   filters = "ensembl_gene_id",
                   values = SearchList,
                   mart = ensembl) 
  row.names(matrix_topx) <- gene.name[,2]
  
  #group_mean<-matrix (nrow=nrow(matrix),ncol=1) #creates one column matrix to contain the means of the control group
  #create matrix for one-way comparison
  #for (j in 1:nrow(matrix)) {
   # group_mean[j,]<-mean(matrix[j,7:12])
    #loop finds the average of the control group and puts them in the matrix 'group mean'
  #}
  #one.way.matrix<-matrix [,1:6] #makes matrix of the experimental group (i.e. exp grp and not control grp)
  #one.way.matrix<-cbind (one.way.matrix, group_mean) #combines the two matricies together
  #one.way.matrix<-one.way.matrix[,1:6]/one.way.matrix[,7] #changes the value of each cell to the count/mean of the control group and then removes the excess column
  ##
  
  
  
  colour<-c(rep("dodgerblue1",6),rep("firebrick",6))
  htmpname<-paste0(subdir[i],"heatmap_of_DEGS_scaled_by_row_",comp[i],".pdf")
  pdf(htmpname)
  heatmap.2(matrix, col=bluered(100), trace="none",scale="row",dendrogram = "row", Colv = FALSE, ColSideColors = colour, sepwidth=c(0.00001,0.00001),cexRow=0.5,cexCol=0.3) # Conv=false for no column dendrogram reordering
  heatmap.2(matrix, col=bluered(100), trace="none",scale="row",Colv = TRUE, ColSideColors = colour, sepwidth=c(0.00001,0.00001),cexRow=0.5,cexCol=0.3) # Conv=True column dendrogram reordered #complete Eculidian clustering
  heatmap.2(matrix_topx, col=bluered(100), trace="none",scale="row",dendrogram = "row", Colv = FALSE, ColSideColors = colour, sepwidth=c(0.00001,0.00001),cexRow=0.5,cexCol=0.3) # Conv=false for no column dendrogram reordering
  heatmap.2(matrix_topx, col=bluered(100), trace="none",scale="row",Colv = TRUE, ColSideColors = colour, sepwidth=c(0.00001,0.00001),cexRow=0.5,cexCol=0.3) # Conv=True column dendrogram reordered #complete Eculidian clustering
  
  #then make FC vs other group heatmap...
  
  
  #heatmap on FC matrix (Which I need to create!) scale="none"

  dev.off()
}  


#rm to remove variables at end of loop to error trap.
```
